# MorphLink: Bridging Cell Morphological Behaviors and Molecular Dynamics in Multi-modal Spatial Omics

**DOI:** 10.1101/2024.08.24.609528

**Authors:** Jing Huang, Chenyang Yuan, Jiahui Jiang, Jianfeng Chen, Sunil S. Badve, Yesim Gokmen-Polar, Rossana L. Segura, Xinmiao Yan, Alexander Lazar, Jianjun Gao, Michael Epstein, Linghua Wang, Jian Hu

## Abstract

Multi-modal spatial omics data are invaluable for exploring complex cellular behaviors in diseases from both morphological and molecular perspectives. Current analytical methods primarily focus on clustering and classification, and do not adequately examine the relationship between cell morphology and molecular dynamics. Here, we present MorphLink, a framework designed to systematically identify disease-related morphological-molecular interplays. MorphLink has been evaluated across a wide array of datasets, showcasing its effectiveness in extracting and linking interpretable morphological features with various molecular measurements in multi-modal spatial omics analyses. These linkages provide a transparent depiction of cellular behaviors that drive transcriptomic heterogeneity and immune diversity across different regions within diseased tissues, such as cancer. Additionally, MorphLink is scalable and robust against cross-sample batch effects, making it an efficient method for integrative spatial omics data analysis across samples, cohorts, and modalities, and enhancing the interpretation of results for large-scale studies.

## Introduction

Histological images are widely used to characterize complex tissue phenotypes related to diseases. Visual inspection of biopsy specimens through histopathology has long been considered as the gold standard for disease diagnosis, as disease associated cells often exhibit distinct morphological changes, such as alterations in size, shape, and the organization of cells and extracellular matrix[1]. These changes in cell morphology and the arrangement of extracellular structures can alter cells’ physical capabilities, affecting their interactions with the environment, adhesion to surfaces, and migration. Complementing the analyses of morphology, high-throughput omics profiles—including mRNA expression, protein abundance, and chromatin accessibility—provide a snapshot of the cells’ functional states and their responses to environmental stimuli. The joint analysis of tissue morphology and molecular dynamics has played a pivotal role in revealing how a cell changes morphology to adapt to its function, identifying disease-related morphology-molecular relationships, and investigating the underlying causes of aberrant cellular behaviors in diseases[2].

Spatial omics techniques represent the latest frontier in high-throughput omics profiling[3]. These techniques measure diverse omics modalities while preserving their native tissue contexts. Key techniques in spatial omics include Spatial Transcriptomics (ST)[4-6], which measures spatial mRNA abundance; spatial proteomics[7, 8], which measures high-plex protein abundance; spatial CITE-seq[9-11], enabling the simultaneous measurement of proteins and the transcriptome; and spatial metabolomics[12] for detailed metabolite profiling. Data from these technologies are often paired with high-resolution hematoxylin and eosin-stained (H&E) images, offering cell morphology information from the same tissue slice. Such multi-modality data provide a valuable resource for linking cell morphology alterations with molecular changes. These linkages are crucial for understanding biological processes from multiple perspectives.

Several pioneering methods have been developed for the joint modeling of molecular information and histology images in spatial omics analysis. For instance, MUSE[13] and SiGra[14] utilize multi-view autoencoders to extract and combine image features with gene expression data; SpaGCN[15] and SpatialGlue[16] leverage undirected weighted graphs to combine molecular and image information; TESLA[17] and iStar[18] impute gene expression data into super-resolution gene images by leveraging high-resolution information provided by histology images. However, these methods are primarily designed for spatial domain detection by clustering and focus mainly on expression data, with histology playing a supporting role. After clustering, a common challenge is determining whether those identified spatial domains are true biological entities or just artifacts resulting from technical noise. This necessitates a post-analysis examination to assess the coherence between histological features and molecular expression within each domain. Currently, no existing methods specifically address the need to bridge studies between morphology and molecular profiles. Efforts to assess the relationship between cell morphology and omics measurements largely depend on manual examination by pathologists[19-21], which are prone to errors and inter-reader variability[22-24]. Therefore, there is a pressing need for methods that can systematically and quantitively examine the morphology-molecular relationships in large tissue sections.

Addressing this need is non-trivial due to the following challenges: First, most analytic methods for spatial omics data rely on well-trained deep neural networks[25-27] to extract hundreds of image features from histology images. Unlike genes and proteins that have clear biological meanings and functions, image features from neural networks are often not transparent, making biological interpretation difficult. Second, training such deep neural networks requires a substantial number of annotated images, leading to a labor-intensive labeling process by pathologists and inconsistent performance across different tissue types. Third, although tools like QuPath[28], CellVit[29], and Hover-Net[30] can extract morphology features from histology images, they primarily focus on nuclear structures and often overlook other non-nuclei structures, such as the extracellular matrix and collagen fibers, which provide complementary insights into the pathological states of tissues. As a result, these tools generate only a limited number of features, which are inadequate for linking morphology with high-dimensional molecular measurements. Lastly, there is a shortage of metrics or methods to quantify the relationship between tissue morphological and molecular measurements within a spatial context.

Here, we introduce a new framework, MorphLink, to address these challenges simultaneously. MorphLink has three primary goals: 1) To extract comprehensive morphological measurements with high interpretability in a label-free manner. 2) To efficiently quantify the relationships between cell morphological and molecular features in a spatial context. 3) To visually demonstrate how cellular behavior changes from both morphological and molecular perspectives. Using diverse types of spatial omics datasets, we showcase MorphLink’s efficiency in extracting and linking various morphological hallmarks with different molecular measurements. These linkages help researchers better understand the biological properties of identified spatial domains and illustrate the mechanisms underlying tumor heterogeneity and immune diversity in multiple cancer types. Given the limited use of spatial omics in routine practice due to cost and accessibility constraints, we strongly believe that the interpretable and biologically relevant morphology features identified by MorphLink will catalyze the advancement of molecularly informed histopathology analysis, paving the way for developing accurate predictive and prognostic risk models based on histology images.

## Results

### Overview

We explain the workflow of MorphLink using ST data as an example, but the method is also applicable for various types of spatial omics analyses. As shown in **Fig. 1a**, MorphLink begins by extracting image patches from the H&E image for measured spots in a ST tissue section. These patches then undergo spatially aware, unsupervised segmentation to generate multiple binary masks, each representing a particular structure. The properties of each mask are summarized to help users identify which cellular/extracellular structure each mask measures, such as nuclei, stroma, and collagen fibers. The distribution of white pixels on each mask demonstrates the proportion and organization of target structures. Summary statistics are calculated as mask-level features to quantify the distribution of these structures in each patch, capturing tissue niche layout heterogeneity. Next, connected component detection is performed within each mask to identify objects such as individual nuclei, stromal aggregates, and fiber bundles. Shape properties such as area, orientation, and solidity are measured to describe the physical attributes of each object. These object-level features complement the mask-level features, which measure tissue organization, by providing insights into sub-cellular textural morphology. As detailed in **Supplementary Table 1**, Morphlink can extract 10 mask-level features and 109 object-level features from each mask. Each H&E image usually has 8-10 masks, resulting in around 1,000 features that can be are easily interpreted with morphological significance.

**Fig. 1.**
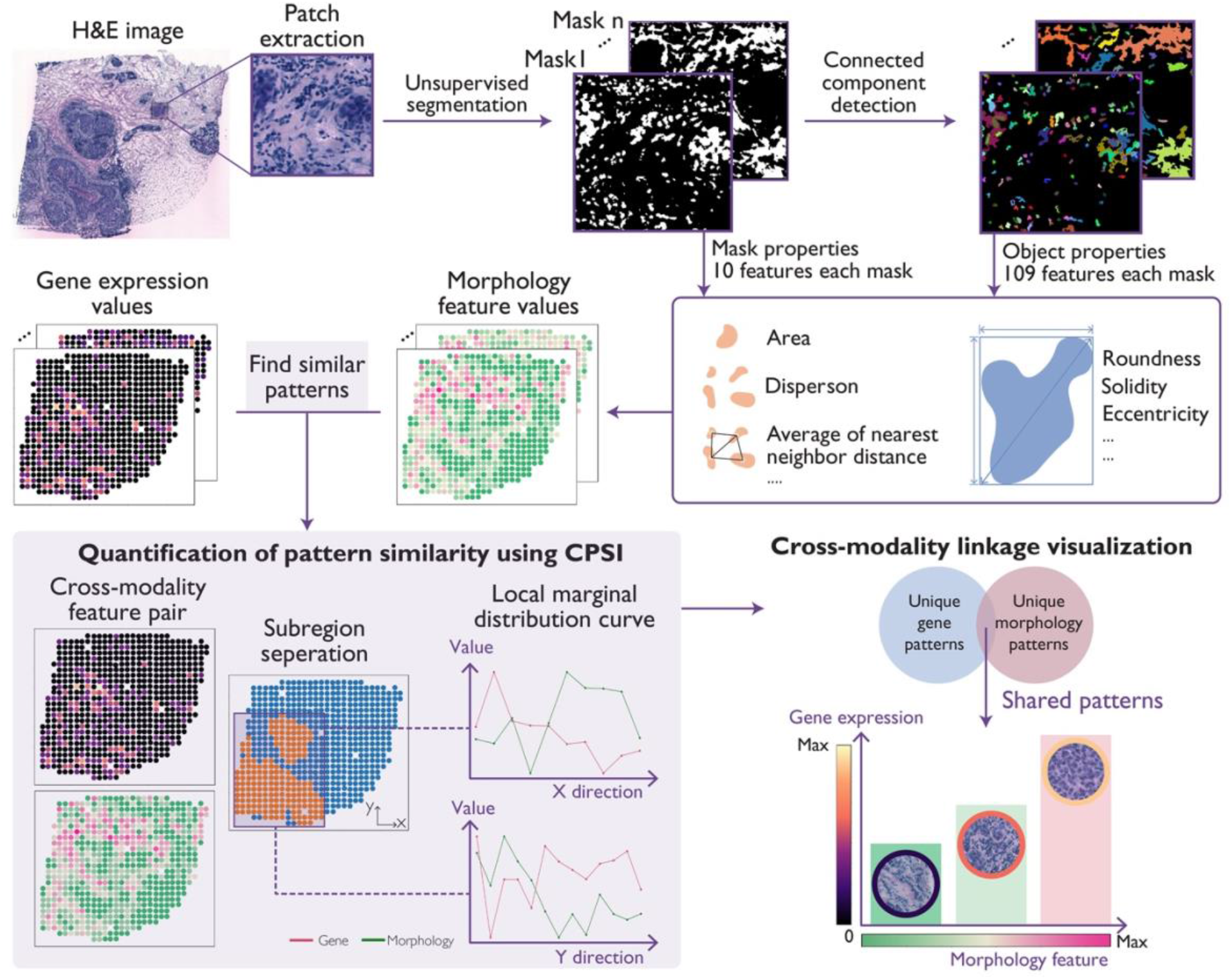
Workflow of MorphLink. **a**. MorphLink starts from extracting image patches from the H&E image, then employs an unsupervised, spatially aware approach to segment each patch to multiple masks. Each mask represents a type of tissue structure, and object detection is further performed. Both mask-level and object-level summary statistics are calculated to generate interpretable image features. **b**. MorphLink quantifies the pattern similarity between morphological and molecular features by initially partitioning the tissue into subregions. Within each subregion, it calculates gradient curves along the X and Y directions to summarize the patterns of both features. The Curve-based Pattern Similarity Index (CPSI) is then computed based on these regional curves. Next, MorphLink samples patches to visually illustrate the dynamic changes in cell morphology and gene expression.

To understand the relationship between tissue morphology and molecular characteristics, it’s essential to identify features of both modalities that share similar distribution patterns. MorphLink introduces a new statistical metric, the Curve-based Pattern Similarity Index (CPSI), as shown in **Fig. 1b**, to quantify the similarity of these patterns between paired features from different modalities. Our observations suggest that many features exhibit strong spatial patterns within specific tissue regions, rather than uniformly across the entire section. To capture such localized patterns when comparing similarity, the calculation of CPSI begins by partitioning the entire tissue section into subregions in a data-driven manner. In each subregion, a 2D feature pattern is broken down into changes along two orthogonal directions. Thus, the spatial pattern of each feature in the subregion is described using marginal curves along the x and y directions.The marginal similarity of a feature pair is quantified by a weighted sum of their curve correlation and absolute difference. The subregion-level pattern similarity is then calculated as a weighted sum of their marginal similarities. Leveraging CPSI, MorphLink is able to identify features with similar spatial patterns across different modalities both locally and globally. Finally, MorphLink selects patches based on their feature values and highlight the measured structures to visually showcase what morphology features and how they change along with gene expression dynamics.

To showcase the effectiveness of MorphLink, we applied it to four bi- or tri-modality spatial omics datasets listed in **Supplementary Table 2**. In each case, MorphLink effectively discerned multiple interpretable morphology features linked to different molecular measurements, underscoring unique cellular behaviors that drive tumor heterogeneity and immune diversity from both morphological and molecular perspectives. Additionally, we extensively evaluated the novel metric CPSI, revealing its ability to detect spatial pattern similarities, outperforming traditional metrics including correlation, structural similarity index measure (SSIM), and root mean squared error (RMSE). MorphLink’s ability to analyze a broad spectrum of spatial omics data and its resilience to batch effects make it an optimal tool for transparent analysis of multi-sample spatial omics data generated from diverse studies.

### Interpretable morphology features extraction

To illustrate MorphLink’s ability to extract interpretable morphological features, we first analyzed a human bladder cancer ST dataset. The H&E image with pathology annotation is shown in **Fig. 2a**, and we focused on the spots enriched with tumor cells. Following the standard ST data analysis pipeline, we first performed spatial clustering to delineate the tumor regions into two distinct subtypes in **Fig. 2b** based on their gene expression profiles. To understand the unique expression pattern of each region, we performed spatially variable gene (SVG) detection to identify genes that show enriched expression pattern in each of the tumor subtype regions. Region 2 stood out with a unique expression pattern, represented by high levels of 977 SVGs including antigen-presenting genes (e.g., *CD74, B2M*, and *TAP1*) and genes associated with tumor proliferation (e.g., *MYCL, MKI67*, and *TUBB*), as illustrated in **Fig. 2c** and **Supplementary Fig. 1**. The elevated expression of these genes suggests that tumor cells in region 2 are more effective at antigen presenting[31, 32] and exhibit faster growth[33-35], creating a more aggressive tumor environment. To show extract interpretable image features that capture tumor heterogeneity, MorphLink began by identifying the key morphological structures in the target region. It employed a spatially aware unsupervised segmentation technique, generating eight masks as depicted in **Supplementary Fig. 2**. The properties of each mask are summarized in **Supplementary Table 3**. Specifically, masks 1 and 2 capture the nuclei and cancer-associated fibroblasts (CAFs), which are the dominant structures in the cancer region, and we use these two masks for downstream illustration. We selected two Visium spots from region 1 and region 2 marked in **Fig 2b** as examples. Visual examination of **Fig. 2d** indicates that spot 2 has larger nuclei and a higher density of CAFs compared to spot 1. MorphLink captures these differences by measuring the interquartile range (IQR) of nuclei solidity in mask 1 (spot 1 = 0.11 vs. spot 2 = 0.18) and the area of CAFs (spot 1 = 3.7 × 10^4^ pixels vs. spot 2 = 7.3 × 10^4^ pixels). The patterns of these two features, as presented in **Fig. 2e**, exhibit elevated values in region 2, distinguishing it from region 1. This example showcases how MorphLink quantifies tissue morphology in an interpretable manner.

**Fig. 2.**
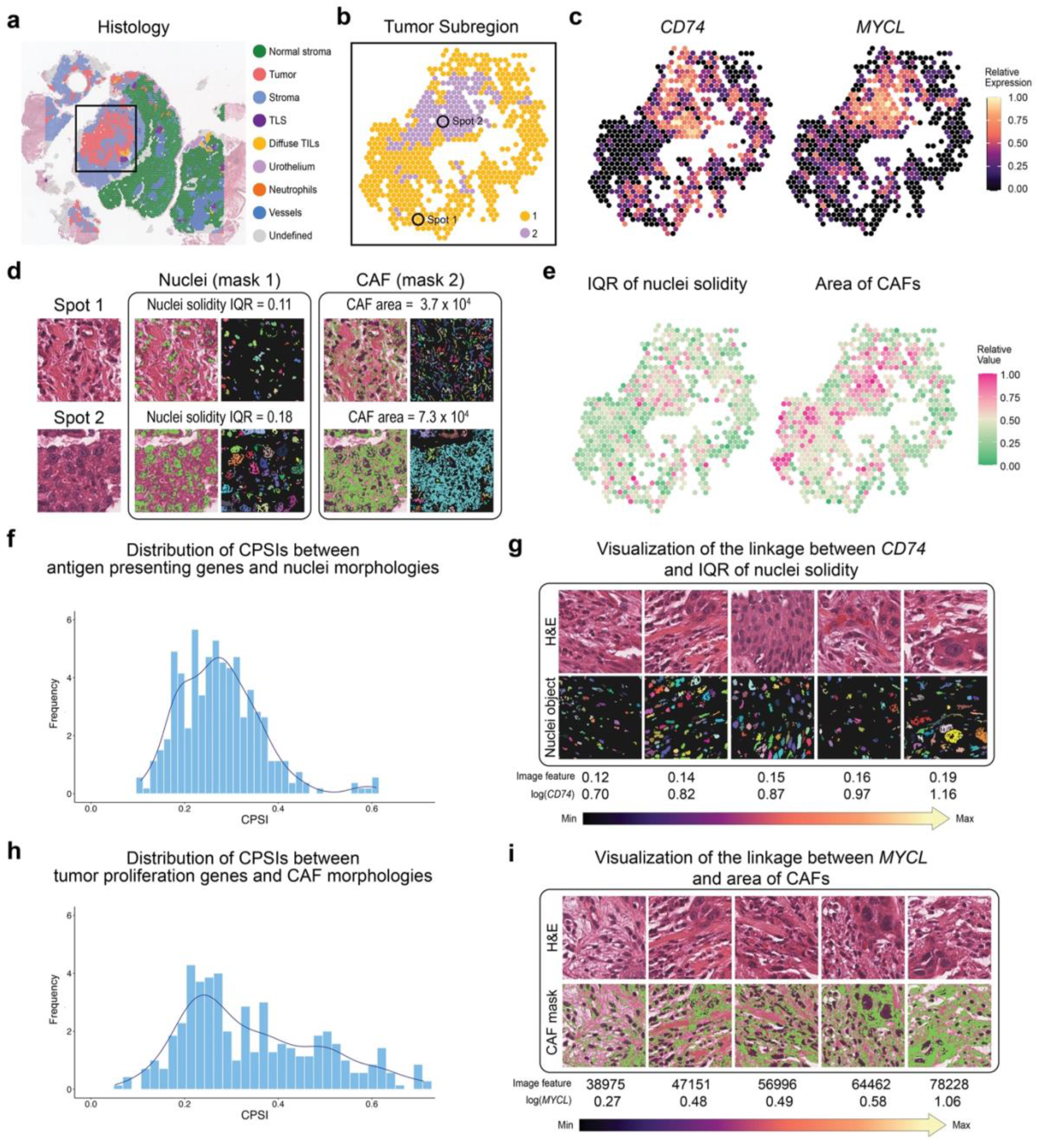
MorphLink associates nuclei and CAF morphology with gene expression to characterize tumor heterogeneity in human bladder cancer. **a**. Pathologists annotations at the spot level overlaid on the H&E image. **b**. The selected tumor region is further divided into two subregions by spatial clustering using gene expression. **c**. The expression patterns of *CD74* and *MYCL* in selected tumor region. **d**. H&E patches of two spots, with two corresponding segmented masks for nuclei and CAF; object detection is subsequently performed within each mask. **e**. Morphological features from MorphLink that quantifies the interquartile range (IQR) of solidity for nuclei and area of CAFs. **f**. Distribution of CPSIs between antigen presenting genes (*CD74, B2M, TAP1*) and nuclei morphology features from MorphLink. **g**. A visual illustration of the linkage between *CD74* expression and IQR of solidity for nuclei in H&E image and detected objects. The values are median image feature values for spot stratified by *CD74*’s expression level, grouped into quantiles from 0 to 1 with step of 0.25. **h**. Distribution of CPSIs between tumor proliferation genes (*MYCL, MKI67, TUBB*) and CAF morphology features from MorphLink. **i**. A visual illustration of linkage between *MYCL* expression and area of CAFs in H&E image and segmented mask. The values are median image feature values for spot stratified by *MYCL*’s expression level, grouped into quantiles from 0 to 1 with step of 0.25.

### Linking nuclei morphology with tumor antigen presentation

Next, we demonstrate MorphLink’s capability to find the linkage between nuclei morphology and molecular characteristics, highlighting cellular behavior heterogeneity within a tumor. SVG analysis has identified a list of genes, as depicted in **Fig 2c** and **Supplementary Fig. 1**, which are involved in antigen presentation and are notably enriched in region 2. Their increased expression suggests that cells in region 2 are more active at the processing and presenting of antigens, potentially making them more recognizable to the immune system compared to region 1 with lower expression of these molecules [36, 37]. To investigate how this biological process influences tumor nuclei, we calculated the CPSIs between antigen-presenting genes and the 119 nuclear morphology features. The results, displayed in **Fig. 2f**, reveal that MorphLink selected a feature consistently demonstrating high CPSIs with all antigen-presenting genes, including *CD74* (CPSI = 0.58), *B2M* (CPSI = 0.56), and *TAP1* (CPSI=0.60). This feature measures the IQR of the nuclei solidity as previously described in **Fig 2e**. Variations in this feature’s value indicate significant shifts in chromatin organization, observable as dark-violet stained structures within nuclei on H&E images. The compact form of DNA can influence nuclear structure and size, correlating with changes in gene expression [38]. To further investigate their relationship, MorphLink uses *CD74* as an example to generate a visual representation. As shown in **Fig. 2g**, with increased expression of *CD74*, the nucleus transitions from dark, dense chromatin to more pronounced clumping of dispersed chromatin, with inconspicuous nucleoli and scant cytoplasm. This transformation, from heterochromatin to euchromatin, typically occurs in regions that are initially transcriptionally inactive. The loosening and opening of chromatin typically marks an increase in transcriptional activity, including the transcription of genes involved in antigen presentation. Previous research has linked certain genetic changes to altered expression of genes that are involved in regulating transcriptional activity and chromatin remodeling[38]. Further analysis revealed an enrichment of these genes in region 2 (**Supplementary Fig. 3**), which is on par with our observations.

### Linking CAF morphology with tumor proliferation

Apart from the tumor cell nuclei, MorphLink can also examine morphology for non-nuclei structures such as stroma. We emphasize the significance of non-nuclear structures by highlighting the tumor proliferation status in region 2. In addition to the antigen-presenting genes mentioned earlier, another set of genes enriched in region 2, including *MYCL* (**Fig. 2c**), *TUBB*, and *MKI67* (**Supplementary Fig. 1**), are cell cycle-related genes that regulate cell growth and proliferation. Overexpression of these genes can lead to the uncontrolled division and growth of tumors and CAFs, a hallmark of an aggressive tumor region[39, 40]. Identifying CAFs on H&E images can be challenging because their nuclei are typically smaller, less dense, and more irregular compared to those of tumor cells and lymphocytes. Furthermore, CAFs are embedded within stroma, a complex mix of extracellular matrix and fibers. This dense stromal background can obscure the subtle characteristics of CAF nuclei, complicating their identification with nucleus-focused image analysis tools. MorphLink is designed to analyze broader morphological features beyond just nuclei, with mask 2 in **Fig. 2d** captures the CAFs, highlighted by their surrounding stroma. By calculating the CPSIs between tumor proliferation genes with 119 morphology features for CAFs (**Fig. 2h**), MorphLink identified a morphology feature (**Fig. 2e)**, the area of the CAF, which has the top CPSIs with genes relevant to proliferation, including *MYCL* (CPSI = 0.637), *MKI67* (CPSI = 0.506) and *TUBB* (CPSI = 0.378). The visual representation in **Fig. 2i** reveals that an increase in *MYCL* expression is directly linked to an expansion of the CAF region surrounding tumor cells, as highlighted in green. This expansion plays a pivotal role in supporting the growth of tumor cells, enhancing their motility, and making them more conducive to infiltration. This CAF-related feature, along with the changes in nuclear chromatin noted in our previous analysis, elucidates altered cell behaviors and highlights tumor heterogeneity in a complementary manner.

### Linking lymphocyte organization with immune diversity

In addition to identifying morphology-molecular linkages highlighting tumor heterogeneity, MorphLink can also capture alterations in cell organization coupled with changes in gene expression related to immune diversity. Lymphocytes in the baldder cancer tissues can be categorized as either part of tertiary lymphoid structures (TLS) [41] or as diffused tumor-infiltrating lymphocytes (TIL), as shown in **Fig 3a**. SVG analysis revealed two genes, *IGHM* and *MS4A1* in **Fig. 3b**, can distinguish these two lymphocyte organizations. Both genes are well-reported for their critical role in lymphocyte infiltration and TLS formation[42]. MorphLink selected the feature in **Fig. 3c** that has highest average CPSIs with *IGHM* (CPSI = 0.483) and *MS4A1* (CPSI = 0.459). The distribution of CPSIs between these two genes and all lymphocyte nuclei features is shown in **Supplementary Figure 4**. This feature measures the largest cluster size of lymphoid nuclei aggregation. Immune cells, which have a high nucleus-to-cytoplasm ratio, are identified in H&E images by the arrangement of their nuclei. Thus, the size of these clusters indicates how closely lymphocytes are grouped together. Notably, this morphology feature’s value in TLS is significantly larger than in diffused TLS, as demonstrated by the boxplot in **Fig. 3d**. A one-sided t-test confirms this distinction with a p-value of 3.24e-09. The visualization provided by MorphLink in **Fig. 3e** also shows that in TLS, a high value of this feature indicates that lymphocytes are forming compact clusters. In contrast, within diffused TIL regions, this feature has a lower value, suggesting a looser, more dispersed distribution of the lymphocytes. The function and interaction of immune cells are influenced by their spatial organization. MorphLink can effectively capture these differences, offering insights into the immune diversity. In conclusion, the examples illustrate MorphLink’s ability to identify morphological features in tumor, CAF, and immune cells, encompassing a broad range of properties including their shape, layout, and organization. These features, closely linked to molecular dynamics, provide a comprehensive analysis of tissue morphology and its functional implications.

**Fig. 3.**
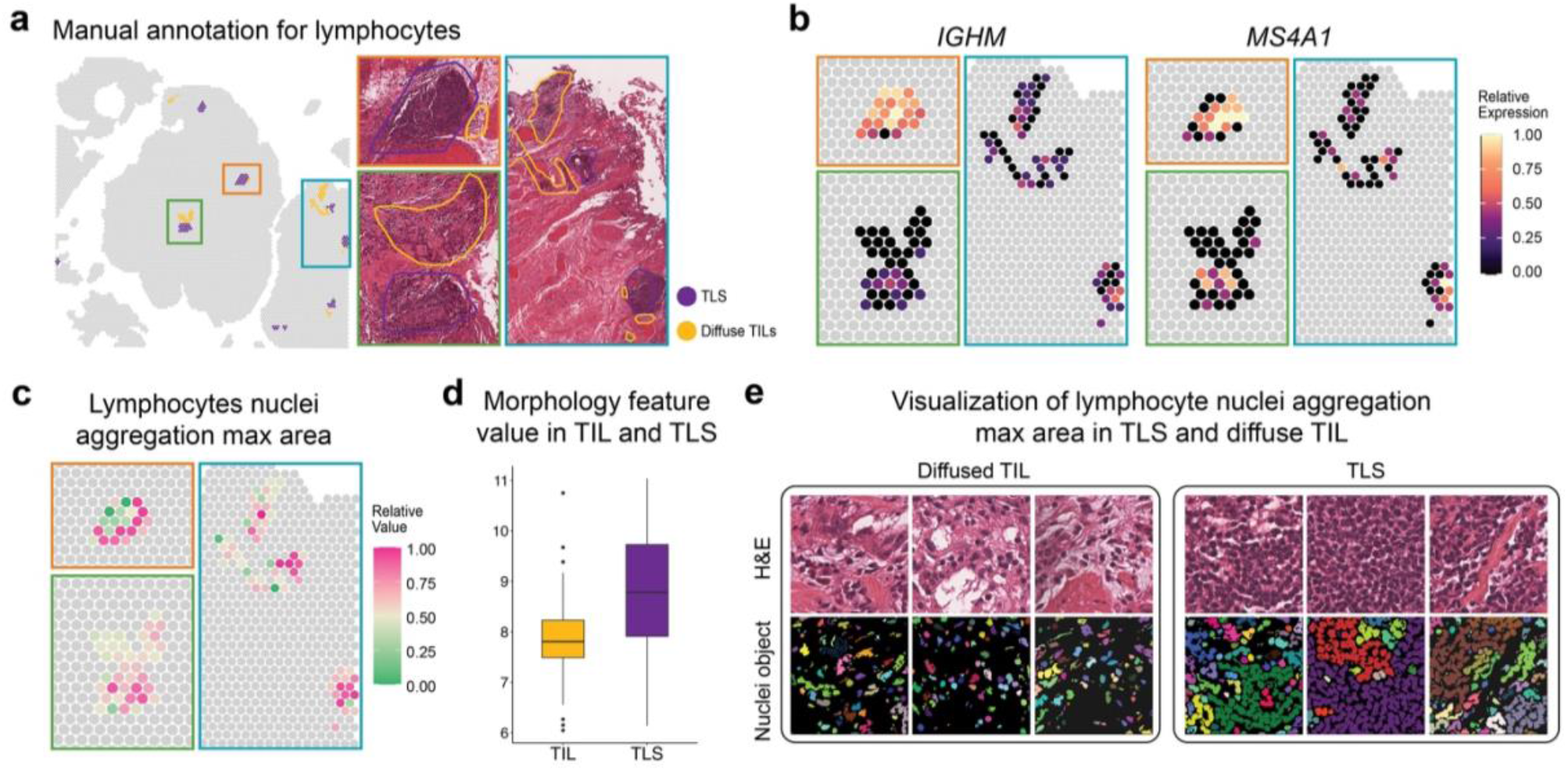
MorphLink associates lymphocyte organization with gene expression to characterize immune diversity in human bladder cancer. **a**. Tertiary lymphoid structures (TLS) and diffuse tumor-infiltrating lymphocytes (TIL) annotation on spot-level and H&E image from the pathologist. **b**. The expression of *IGHM*, and *MS4A1* at lymphocyte enriched regions. **c**. A morphological feature that measures the maximum area of lymphoid nuclei shows high CPSIs with all TLS enriched genes. **d**. Boxplot of the morphology feature value in TIL(n=73) and TLS (n=88). The lower and upper hinges correspond to the first and third quartiles, and the center refers to the median value. The upper (lower) whiskers extend from the hinge to the largest (smallest) value no further (at most) than 1.5 × interquartile range from the hinge. Data beyond the end of the whiskers are plotted individually. **e**. A visual illustration depicting how the organize of limnophytes differs between TLS and diffused TIL in H&E image and detected objects.

### Application to multi-sample ST data

As spatial omics technology becomes more affordable, an increasing number of studies are producing spatial omics data from experiments designed with cohort-level sample sizes[43-45]. Therefore, it’s crucial for newly developed methods to be scalable for multi-sample analysis. We further illustrate MorphLink’s ability to process data from multiple samples by identifying consistent morphology-molecular linkages across various samples within a cohort. We analyzed a HER2+ human breast cancer dataset[43] collected from eight patients, but we focused on tissue sections from patients A, H, and G, as these sections featured both invasive cancer and regions of ductal carcinoma in situ (DCIS), as shown in **Fig. 4a**. Our initial approach involved differential gene expression analysis aimed at identifying genes showing variations across different tumor subregions. This analysis revealed that certain collagen genes, such as *COL1A1, COL1A2*, and *POSTN* depicted in **Fig. 4b** and **Supplementary Figure 5**, exhibited higher expression in invasive cancer regions compared to DCIS across all samples. MorphLink selected the image feature with highest average CPSI to *COL1A1* across three samples, showcased in **Fig 4c**. The distributions of CPSIs for each sample is shown in **Supplementary Figure 6**. From the mask properties listed in **Supplementary Table 4**, we can determine that this mask (mask 2) captured the stroma region in H&E images, and this feature quantifies the spatial continuity of stroma by measuring the IQR of distances between stroma pixels. It consistently exhibits higher values in DCIS than in invasive cancer regions as illustrated in the boxplot in **Fig. 4d**. One-sided t-tests reveal p-values of 0.01, 0.07, and 2.3e-07 for samples A1, G2, and H1, respectively. The examples provided by MorphLink in **Fig. 4e** illustrate that in invasive cancer, the stroma region not only occupies a larger proportion but also exhibits greater integration, filling more space between tumor cells compared to DCIS. The increased area and expanded presence of the stroma region are indicative of a microenvironment conducive to tumor cell growth and invasion [46]. This observation points to a more aggressive and actively progressing state in the invasive tumor region compared to DCIS, highlighting the effectiveness of MorphLink in differentiating between these two stages of tumor development across multiple samples. Additionally, compared to deep neural network features, features extracted by MorphLink exhibit superior resilience to batch effects arising from variations in staining across different rounds (**Supplement Note 1**), making them better suited for cohort-level ST data analysis.

**Fig. 4.**
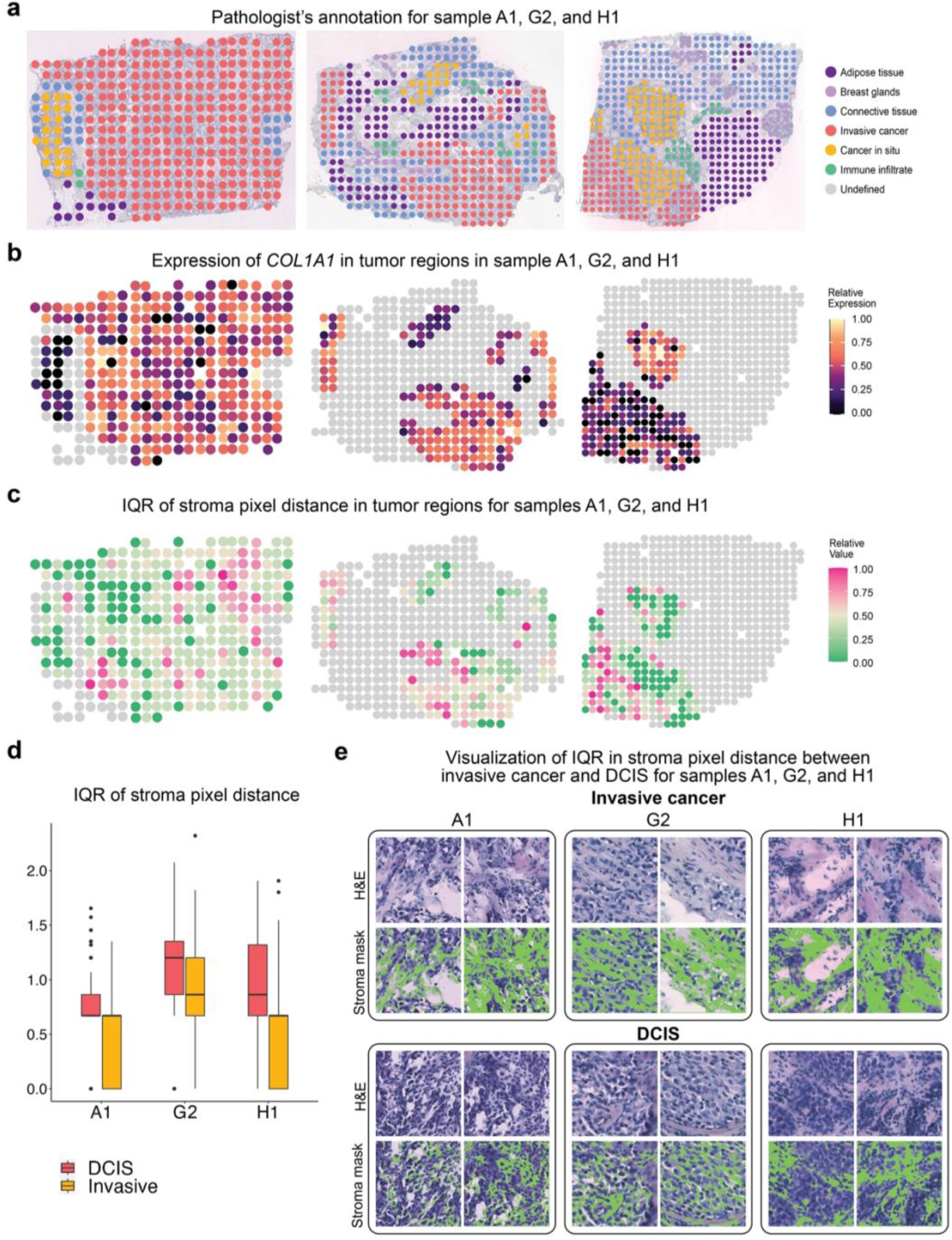
MorphLink identified consistent morphology-molecular relationships in a multi-sample human breast cancer dataset. **a**. The H&E images of tissue sections from patients A, G, and H with manual annotation. **b**. Expression of *COL1A1* in tumor regions across tissue sections of A1, G2, and H1. **c**. A morphological feature that quantifies the density and spatial continuity of the stroma pixels consistently exhibits the highest CPSI with *COL1A1* across all sections. **d**. Boxplot of the morphology feature value in DCIS and invasive cancer region in A1 (n=282, 21), G2 (n=140, 20), and H2(n=90, 97). Boxplot hinges, median and whiskers are defined the same as in 2d. **e**. A visual illustration show that higher image feature value indicates that the stroma region exhibits a greater density and a more expanded presence between tumor cells in the invasive cancer region than in the cancer in situ region in sample A1, G2, H1.

### Illustration of CPSI

An important step in MorphLink is to quantify spatial pattern similarity using our proposed metric, CPSI. CPSI is a decimal value ranging from -1 to 1, where a value of 1 signifies perfect similarity and a value of - 1 indicates perfect dissimilarity. To illustrate how CPSI works, we analyzed zebrafish melanoma Visium dataset[47], with H&E image shown in **Fig. 5a**. This H&E image has a strong artifact of discrepancies in blurriness, which is highlighted by the red dash line. The annotations from the original study in **Fig. 5b** indicate that the tissue can be largely categorized into three areas: melanoma, normal muscle, and tumor/muscle interface. We specifically selected one gene, *RPL15*, and image feature 1 from MorphLink, both of which exhibit strong spatial patterns, for illustration. *RPL15* is ribosomal subunit coding gene and its elevated expression is associated with abnormal ribosome biogenesis in cancer. As depicted in **Fig. 5c**, *RPL15* it is unsurprisingly highly expressed in melanoma cells. Intriguingly, its expression in muscle cells also increases progressively as they approach the melanoma region, indicating that muscles closer to the tumor experience a greater burden, induce stress responses, and lead to morphological changes. On the other hand, image feature 1 from mask 6, which measures the minium extent of the inter-cellular space (**Supplementary Table 5**), also shows higher values in muscle cells closer to melanoma. This indicates that these cells have contracted, resulting in larger intercellular spaces as an adaptation to the tumor burden (**Supplementary Figure 7**). In the melanoma region, image feature 1 has a distinct pattern compared to *RPL15* as it only measures muscle morphology and shows very low values in melanoma. However, their patterns in muscle region are similar as both describe changes in muscle cells adapted to the tumor burden. To capture such local pattern coherence, the initial step in the CPSI calculation is to automatically separate the tissue into subregions (detailed in the Methods). Morphlink automatically separates the whole slide into muscle regions, including both normal and interface areas, and melanoma regions. Within each region, MorphLink uses a curve to summarize feature marginal distribution in two orthogonal directions, i.e., x and y. Then, the curved-based correlation and difference were calculated for each sub region. As shown in **Fig. 5e** and **f**, the curves in the muscle region along the y-axis demonstrate a high correlation (0.68) and a low difference (0.15), reflecting the increasing trend for both *RPL15* and image feature 1 as muscle cells approach the melanoma. Conversely, in the melanoma region, the curves in both the x and y directions (**Fig. 5g**) show relatively lower correlations (0.4, 0.13) and higher differences (0.26, 0.32), indicating distinct patterns in this area. For comparative purposes, we included summary curves for the entire section in **Fig. 5h**. Without subregion separation, the coherent patterns in the muscle region would be missed, as evidenced by globally low correlations (−0.19, -0.11) and high differences (0.3, 0.34) in both directions.

**Fig. 5.**
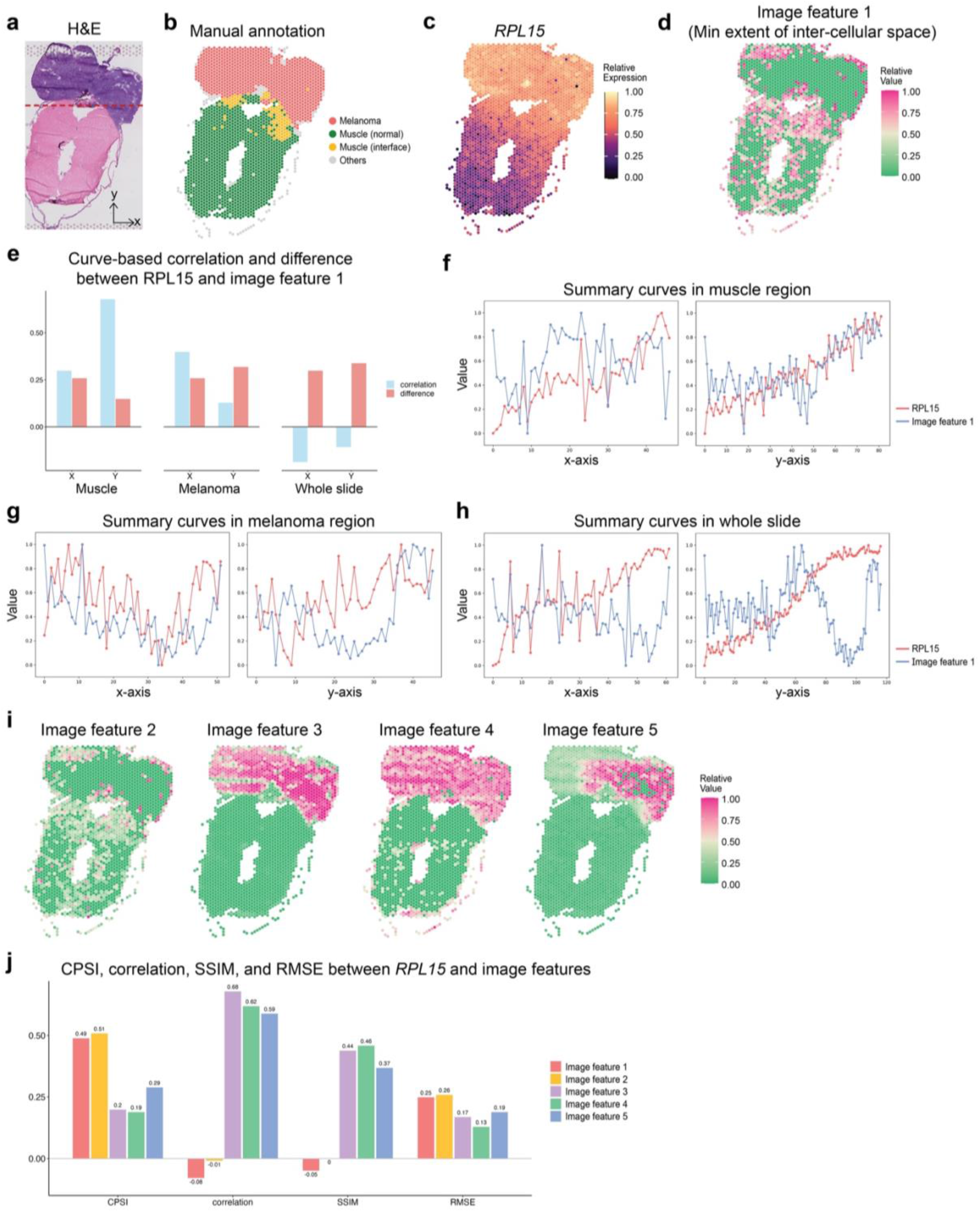
Comparasion of CPSI with other metrics in the zebrafish melanoma dataset. **a**. The H&E image of the tissue section, with the blurred region outlined by a red dashed line. **b**. Manual annotations of the tissue section from the original study. **c**. The spatial expression patterns of *RPL15*. **d**. The spatial pattern of image feature 1. **e**. Barplot of curve-based correlation and difference between *RPL15* and image feature 1 along X and Y directions in subregion 0, subregion 1, and the whole region. **f - h**. Summary curves of *RPL15* and Image feature 1 along two orthogonal X and Y directions in muscle region, melanoma, and the whole slide. **i**. The expression patterns of four additional selected image features: image feature 2 (minimum solidity of inter-cellular space), image feature 3 (proportion of melanoma area), image feature 4 (standard deviation of melanoma pixel distance), and image feature 5 (maximum minor axis length of blurred fiber bundles). **j**. Barplot for CPSI, correlation, Structural Similarity Index (SSIM) and Root Mean Square Error (RMSE) calculated between *RPL15* and image features.

### Evaluation of CPSI

To showcase the effectiveness of CPSI in quantifying morphology and molecular feature similarities, we compared CPSI with other commonly used metrics in the task of pattern similarity quantification, including correlation, SSIM, and RMSE. In addition to image feature 1, we included four additional image features (**Fig. 5i**) in comparison, representing many of the frequent patterns we have observed. Image feature 2 measures the minium solidity of intercellular regions, displaying a pattern similar to image feature 1, but with different magnitudes. Both features are crucial indicators of muscle tissue heterogeneity due to tumor burden and share local patterns similar to *RPL15*. In contrast, image features 3 and 4 from mask 1 are specific to melanoma morphology and do not reflect muscle heterogeneity. Additionally, image feature 5, which affected by the blurriness of the melanoma region, is considered a noise feature resulting from artificial effects. Unlike features 1 and 2, features 3, 4, 5 do not correspond to the patterns observed in *RPL15*. The barplot in **Fig. 5j** compared different metrics between *RPL15* and these image features. Among all the 5 image features, features 1 and 2 have the highest CPSIs, underscoring its capability to efficiently identify coherent spatial patterns. For correlation and SSIM, features 3, 4 and 5 all get the highest values due to their enrichment in the melanoma region. This result highlights the limitation of correlation and SSIM, which tend to capture global patterns but overlook local gradients. Additionally, all five features exhibited low RMSE values, suggesting that RMSE is insufficient for capturing pattern similarity. We also simulated multiple features with distinct spatial patterns to compare the performance of CPSI with other metrics. The simulation demonstrates that CPSI better captures pattern similarity compared to other metrics (**Supplementary Note 2**). Compared to the three competing metrics, CPSI’s superiority is attributed to two key factors: 1) Its ability to automatically separate a tissue into subregions, and 2) the high efficiency of its curve summarization method to capture spatial patterns. These advantages make CPSI a better metric to quantify multi-modal feature similarity in spatial omics data.

### MorphLink is robust in the presence of image artifacts

Deep learning methods for image analysis, such as ResNet[48] and HIPT[25], are often highly sensitive to artifacts, impacting their performance in tasks like clustering. In contrast, MorphLink demonstrates greater robustness to such artifacts. For example, in our analysis of a zebrafish dataset, we noted significant discrepancies in blurriness across the H&E images, marked by a red dashed line in **Fig. 5a**. To demonstrate how this artifact affects downstream analysis, we conducted Louvain’s clustering using image features derived from HIPT and MorphLink separately and the results are shown in **Fig. 6a**. Notably, the domains detected by HIPT features display pronounced artificial effects as the blurred region is identified as cluster 1. By contrast, domains detected by MorphLink features show less impact from blurriness, attributable to the robustness of its interpretable features compared to those generated by deep neural networks. Besides the melanoma region, the muscle region is less affected by artifacts.

**Fig. 6.**
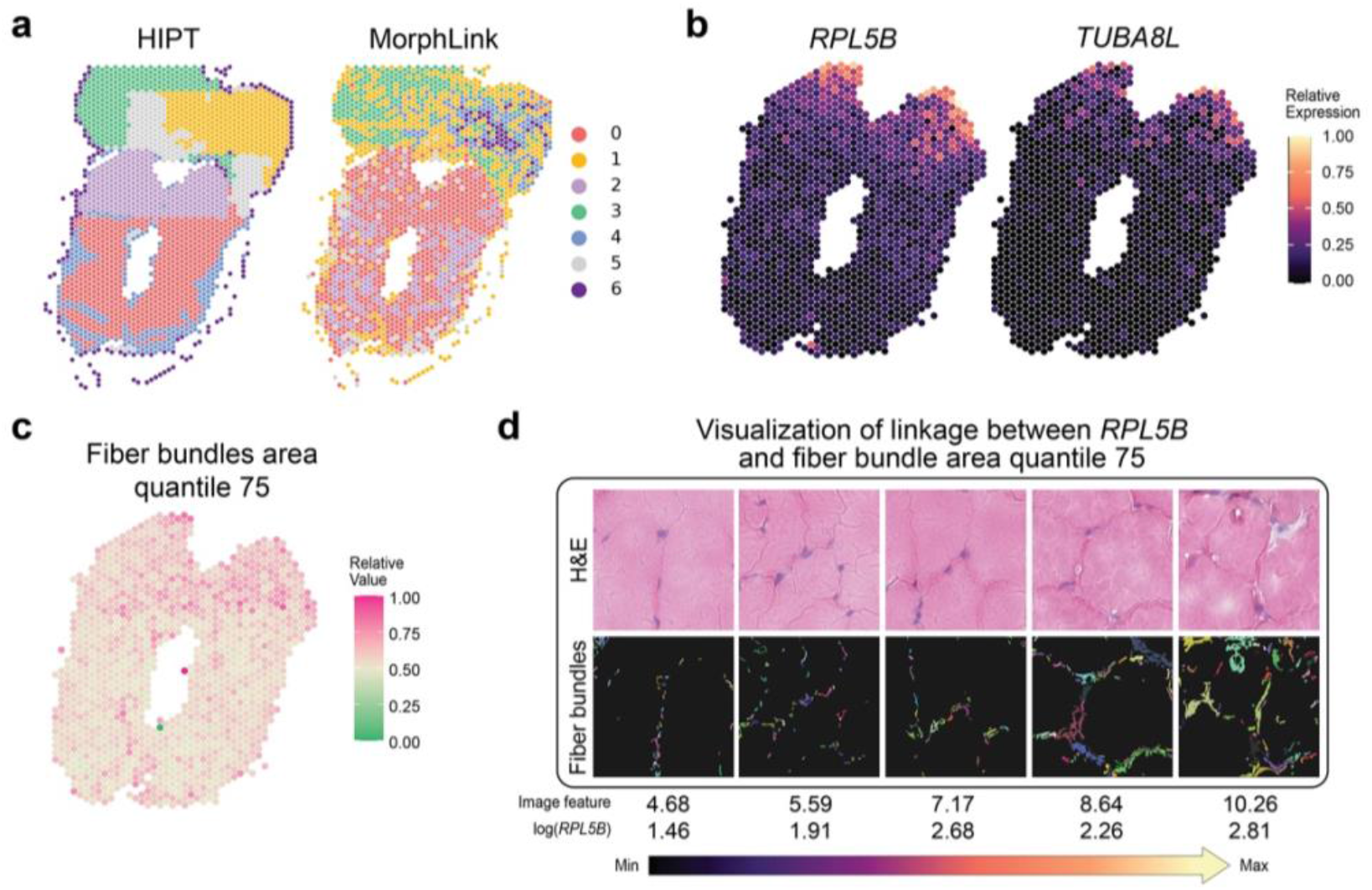
MorphLink characterizes the morphology-molecular relationship at the tumor/muscle interface. **a**. Spatial domains detected using image features derived from HIPT and MorphLink. **b**. Expression patterns of *TUBA8L* and *RPL5B* in the muscle region. **c**. A morphological feature that quantifies the 75^th^ quantile of fiber bundle area shows the highest CPSIs with *RPL5B*(0.757) *and TUBA8L*(0.743). **d**. A visual illustration of muscle fascicle size changes associated with *RPL5B* expression in H&E image, segmented mask, and detected objects. The values are median image feature values for spot stratified by *RPL5Bs* expression level, grouped into quantiles from 0 to 1 with step of 0.25.

We next demonstrate how MorphLink can identify molecular-morphology linkages that distinguish between normal muscle and muscle cells adjacent to the tumor. As reported in the original study, muscle cells near the tumor exhibit distinct gene expression patterns with some example genes shown in **Fig. 6b** and **Supplementary Fig. 8**. We applied MorphLink to all genes enriched in the interface and identified an image feature that has the highest average CPSI (0.73) with all these genes (**Supplementary Fig. 9**). Illustrated in **Fig. 6c** and **Supplementary Table 5**, this feature derived from mask 4 measured the area of fiber bundles at the muscle boundary. Using *RPL5B* as an example, as its expression elevates, the muscle exhibits signs of cachexia at the adhesion sites, characterized by muscle atrophy due to nutrients being consumed by tumor cells[49]. Consequently, the collagen degradation leads to reduction in muscle fascicle size, resulting in more pronounced boundaries in the H&E staining, shown in **Fig. 6d**. This example showcases MorphLink’s capability to detect local morphology-molecular linkage even in datasets with poor quality H&E image, including areas with low resolution.

## Discussion

In this paper, we introduced MorphLink, the very first method designed to extract, interpret and quantify tissue morphology changes associated with molecular profiles using spatial omics data. MorphLink has undergone extensive evaluations across multiple datasets, covering various species tissues, and spatial omics platforms. These results have proven its efficacy in identifying potential morphological markers that indicate molecular alternations leading to tumor heterogeneity and immune diversity. A central feature of MorphLink is CPSI, a novel metric developed for quantifying spatial pattern similarity. CPSI has consistently outperformed traditional metrics such as correlation, SSIM, and RMSE, showcasing its ability to identify features with spatial coherence across multiple modalities.

Compared to existing methods that rely on ‘black box’ modeling, MorphLink surpasses them by providing a higher level of interpretability in spatial omics data analysis. By visually and quantitatively demonstrating the relationship between cellular morphological changes and molecular features, MorphLink allows researchers to clearly distinguish authentic biological patterns in morphology from artifacts, greatly enhancing analytical transparency. Additionally, its label-free nature and robustness against batch effects make MorphLink a superior choice for across sample and cohort data integration. A limitation of MorphLink is that it mainly focuses on capturing local tissue morphology, in contrast to deep neural network features that capture global tissue structures. Therefore, image features from MorphLink may have limitations in clustering tasks that require the identification of global patterns.

In addition to bi-modality spatial transcriptomics data, While this study primarily showcases MorphLink’s application in correlating gene expression with cell morphology in H&E-stained images, MorphLink can also be applied to tri-modality spatial omics data to establish trimodal linkages among protein abundance, gene expression, and morphology, as demonstrated in **Supplementary Note 3**. Beyond H&E stains, MorphLink can be applied to other histological staining techniques, such as Trichrome, Periodic Acid-Schiff (PAS), and Wright’s Stain, which highlight different cellular structures and components. MorphLink’s combination of versatility, interpretability, and scalability renders it an essential tool for the integration of morphology and molecular profiles in multi-sample spatial omics studies.

## Acknowledgements

J.Hu. was fully supported by the start-up funds from the Department of Human Genetics, School of Medicine at Emory University. L.W. was supported in part by the NIH/NCI grants R01CA266280, U24CA274274, The Break Through Cancer Foundation, and the University of Texas MD Anderson Cancer Center Institutional Research Grant (IRG) Program. We thank N. Ma for the workflow figure design and generation.

## Author contributions

This study was conceived of and led by J.Hu.. J.Hu and J.Huang designed the model and algorithm. J.Huang implemented the MorphLink software and led the analysis with evaluation input from S.S.B., Y.G.P., L.W., and M.E.. J.C., J.G., X.Y., R.L.S., and A.L. generated and annotated the human bladder tumor data. C.Y. and J.J. contributed to figure generation. J.Hu and J.Huang wrote the paper with feedback from all other coauthors.

## Methods

### Input data and preprocessing

MorphLink is designed to accommodate a wide range of spatial omics and imaging data from various techniques as inputs. For molecular measurements, including mRNA, protein, or chromatin accessibility, MorphLink is capable of automatically handling raw data and performing all necessary preprocessing steps. To extract morphological features, MorphLink can handle raw histology images collected from different types of staining technologies, including Hematoxylin and Eosin (H&E) stained, Trichrome stained, microscopy, and immunofluorescence imaging.

We demonstrate the method using ST as an example. The spatial gene expression data consist of an *N* × *G* expression matrix with *N* spots and *G* genes, along with the (*x, y*) coordinates to record the 2-dimensional location of each spot. For gene expression and protein abundance, the expression matrix is normalized so that the abundance of each gene/protein in a given spot is divided by the total abundance across all genes/proteins in that spot, multiplied by 10,000, and then transformed to a plus one natural log scale. MorphLink can directly accept raw, high-resolution images and conduct image feature extraction. For multi-image analysis, stain normalization using “staintools” is recommended to mitigate the batch effect across different staining runs.

### Interpretable image feature extraction using unsupervised segmentation

#### Spatially aware patch segmentation

Given the typically higher resolution of histology images (pixel-level) compared to the corresponding omics data (spot-level), it is crucial to extract histological features for each spot to enable the spatial alignment of image and omics features. For each spot *s* in the spatial omics data, its location is represented by the 2-dimensional coordinates (*x*_*s*_, *y*_*s*_). Centered on the coordinates of spot *s*, MorphLink extracts a square patch with a width of *m*. The patch size varies with datasets generated from different techniques, and MorphLink includes a built-in function to help users determine the appropriate patch size. The patch sizes for various datasets are listed in **Supplementary Table 2**. Following this, we obtain a set of patches for all spots stored as a matrix with dimensions of (*N, m, m*, 3), where *N* is the total number of spots. MorphLink then employs a spatially aware image segmentation method to segment each patch into multiple masks. Specifically, for each pixel *ν* in a patch, MorphLink start by extracting its three-channel color values (*r*_*p*_, *g*_*p*_, *b*_*p*_). Taking these values as features, MorphLink applies K-means clustering to divide the pixels into *k* clusters {*c*1, *c*2, …, *ck*}, with (*k* = 10 as default). This clustering step separates pixels based solely based on color value without considering spatial dependence, which may result in non-cohesive, fragile separation. To improve the spatial contiguity of separated clusters, MorphLink uses a convolution filter to refine the cluster assignments. For pixel *ν* assigned to cluster *c*(*ν*) we examine its nearest eight pixels {*u*_1_, *u*_2_, …, *u*_8_} and examine their cluster assignments, denoted as {*c*(*u*_1_), *c*(*u*_2_), …, *c*(*u*_8_)}. We then update the cluster assignment of pixel *ν* by:

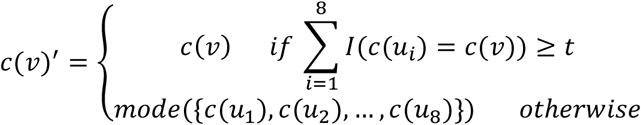

*t* is a parameter set to control the integrity of the clusters, with a default value of 4. This step enhances the spatial continuity of clusters by reassigning a pixel’s cluster when most of its neighbors belong to a different cluster. This refinement process can be performed iteratively until more than 95% of the pixel cluster assignments remain unchanged. Although the pixel-level refinement can enhance the integrity of the segmentation, the initial total number of clusters *k* can be arbitrary and may result in clusters that represent non-cohesive artifacts in the image. Therefore, MorphLink’s next step is to merge these clusters based on their color. For any two clusters *i, j* ∈ {1, 2, … *k*}, we first calculate their color representative as the median for the Red, Green, Blue (RGB) channel separately. Next, we define the color distance between these two clusters as:

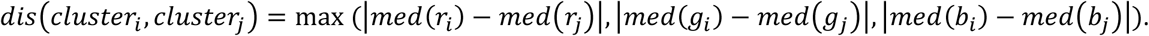

*dis*(*cluster*_*i*_, *cluster*_*j*_) = max (|*med*(*r*_*i*_) − *med*(*r*_*j*_)|, |*med*(*g*_*i*_) − *med*(*g*_*j*_)|, |*med*(*b*_*i*_) − *med*(*b*_*j*_)|). If the distance between two clusters is less than *α*, a predefined threshold, we proceed to merge the two clusters. This approach allows for the generation of a varying number of clusters, customized for different tissue types, ensuring substantial color differences between distinct clusters. Such customization improves the segmentation’s ability to recognize hallmark structures across various tissues. An *α* value of 30 has been consistently effective for segmenting H&E-stained images from diverse tissue types. Subsequently, each merged cluster is converted into a tissue mask to represent a unique biological structure.

#### Mask matching across-patches

The segmentation process is performed within each patch for two primary reasons. First, it allows for parallel processing and thus can be both time and memory efficient. More importantly, segmenting at the patch level enables focused analysis on local tissue morphology, in contrast to the whole slide imaging segmentation, which typically emphasizes large-scale regional distinctions, such as tumor versus non-tumor areas, but overlooks detailed local cellular morphology. This focus on the finer details of local structure is a key distinction between the interpretable features from MorphLink and features from deep learning models.

Given that the initial segmentation is performed independently on each patch, identifying shared clusters across patches is crucial to establish common measurements for consistent tissue hallmarks. Beginning with two patches, labeled *a* and *b*, containing *k*_*a*_and *k*_*b*_ clusters respectively, MorphLink examines each pair of clusters *i, j* across the patches, where *i* ∈ {1, 2, …, *k*_*a*_} and *j* ∈ {1, 2, …, *k*_*b*_}. Utilizing the same methodology as in the cluster merging phase for determining color representativeness, we calculate the color distance *dis*(*cluster*_*i*_, *cluster*_*j*_) between the two clusters. If this distance is less than a threshold of *α, cluster*_*i*_ and *cluster*_*j*_ are considered to represent the same tissue structure. This matching process is applied to all pairs of clusters between patches *a* and *b*. Given that the color differences among clusters within a patch exceed *α*, a single cluster from one patch cannot match multiple clusters from another patch, avoiding any ties. Following this mapping, we end up with *k*_*a*+*b*_ shared clusters, where *k*_*a*+*b*_ ≤ min (*k*_*a*_, *k*_*b*_). These *k*_*a*+*b*_ clusters then serve as the basis for mapping additional patches. By applying this matching process and incorporating one more patch at a time, this approach facilitates the identification of common clusters across all patches, thereby enabling consistent measurements of identical tissue structures across the whole tissue section

#### Interpretable feature extraction

For each mask of a patch, we extract two sets of features: mask-level features and object-level features. Mask-level features aim to quantify the distribution of white pixels in each mask. Additionally, we perform a distance transformation on both the while and black pixels in each mask and utilize summary statistics to characterize their distributions. To extract object-level features, we perform connected component detection on each mask of a patch using the Spaghetti algorithm. This procedure identifies individual objects within the masks, such as single nuclei or stromal aggregation. Following the detection of these objects, we measure the shape properties for each object within a patch mask. Given the numerous objects detected from each patch, we compile summary statistics for each property to obtain single value measurements. These measurements include mean, median, interquartile range (IQR), standard deviation, and quantiles (ranging from 0 to 1 in steps of 0.25). A comprehensive description of extracted image features is listed in **Supplementary Table 1**.

#### Selective-Log transformation

The interpretable image features vary widely in scales and distributions. For instance, features quantifying the area of masks might range from 0 to 10^4^ pixels, whereas the solidity of objects fluctuates between 0 and 1. Inspired by the preprocessing steps used in gene expression data, MorphLink applies a selective log transformation to morphology features to enhance their normality and informativity. For any given feature, we initially adjust it to a non-negative range by subtracting its minimum value. Following this, we apply a plus-one log transformation. We then assess the standard deviation of this feature before and after this log transformation to decide whether the transformation should be retained. Specifically, for a feature *f*, we first normalize it to the [0, 1] range and then compute its standard deviation as *std*_*f*_. Alternatively, we apply a plus-one log transformation on the feature values and then normalize the logged values back to the [0, 1] range. The standard deviation is recalculated as *std*_log (*f*+1)_. If *std*_log (*f*+1)_>*std*_f_, indicating increased variance, we proceed with the plus-one log transformation for that feature. This transformation makes the distribution of image features similar to that of molecular measurements, enabling further comparison.

### Similarity measurement using CPSI

#### Sub region division

Our observations have revealed that many features display strong spatial pattern similarities within specific tissue subregions, rather than uniformly across the entire section. Quantifying these patterns on a whole-section scale could potentially dilute these localized patterns. Hence, our approach with MorphLink begins by partitioning the entire tissue section into subregions. This separation ensures that gene expression and image features overall share a higher degree of spatial pattern similarity within each subregion than across subregions. To achieve this separation, we first perform spatial domain detection through Louvain’s clustering on image features and gene expression data separately. Then the spots are divided into *I* sets by image features, represented as {*s*_11_, *s*_12_, …, *s*_1*I*_}, and into *J* sets by gene expression, represented as {*s*_21_, *s*_22_, …, *s*_2*J*_}. For any cluster pairs with *i* ∈ {1, 2, …, *I*} *and j* ∈ {1, 2, …, *J*}, if

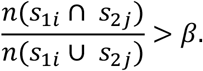

MorphLink proceeds to merge these clusters. To efficiently manage multiple clusters with overlaps that need combination, we sort the pairs of clusters by their overlap ratios in descending order, beginning the merge process with the highest ratio, and evenly results in *K* subregions. By default, the threshold value for *β* is set at 0.2, facilitating a strategic balance in the integration of clusters.

#### Calculating marginal curves

For a feature *f* within a subregion, MorphLink first normalizes its value to [0, 1], and then generates two curves to capture the spatial pattern’s gradient changes along two orthogonal directions, i.e., X and Y. To analyze the feature gradient change along X direction, we first apply a window size of *l* to divide the subregion into *t*_*x*_ intervals of equal length, calculated as:

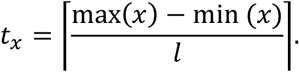

Within each interval, we calculate the median feature value of the spots that fall into it. This process produces a vector of length *t*_*x*_ representing marginal distribution along the x-axis, denoted as

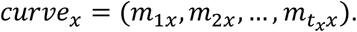

The same procedure is applied to the Y-axis to generate a curve vector of length *t*_*y*_. The window size *l* is the only parameter in this calculation. Rather than directly setting the value of *l*, we control the value of min(*t*_*x*_, *t*_*y*_). More number of intervals, the more detailed the pattern captured by the summary curve.

We set value of *l* so that the min(*t*_*x*_, *t*_*y*_) equals 100, which has consistently yielded good performance across all the analyzed datasets.

#### Quantification of curve-based similarity

For any two features *f*1 and *f*2, the regional pattern similarity can be calculated using the marginal curves as:

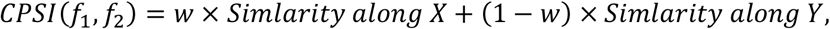

where

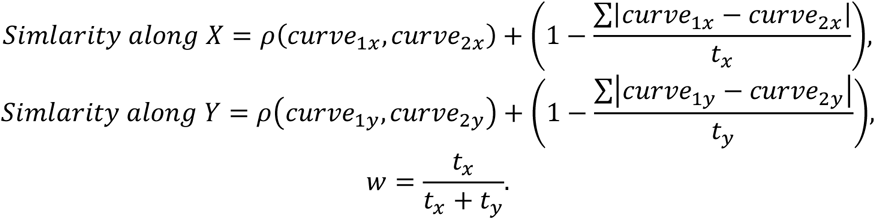

Next, the global CPSI can be derived by summation of local CPSIs a weighed by the size of each subregion:

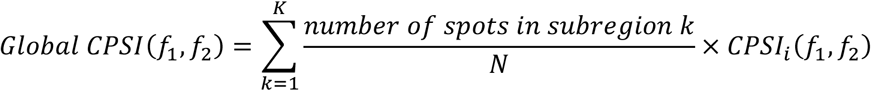

### Sample selection for virtual demonstration

For a pair of morphology feature *f* and gene expression *g* that have a high CPSI, MorphLink provides a visual demonstration of how tissue morphology changes with alterations in gene expression. To accomplish this, MorphLink initially stratifies all the spots into 5 groups based on the quantiles of their *g* expression. Subsequently, it focuses on the tissue masks where feature *f* is measured. The process begins with sorting all the patches in each group based on the area of the target region in each patch, and then selecting those patches whose mask area falls within the 25th and 75th quantiles. This step ensures that the patches chosen for display are proportionally representative of the measured structure. Finally, MorphLink selects the patches with the median *f* value within each group as a representative sample, providing a clear visualization of the relationship between *f* and *g*.

## Data availability

We analyzed three spatial transcriptomics data and one spatial CITE-seq data. Publicly available data were acquired from the following websites or accession numbers: (1) human bladder tumor 10x Visium data (https://www.ncbi.nlm.nih.gov/geo/query/acc.cgi?acc=GSE246011), the following secure token has been created to allow review of record GSE246011 while it remains in private status: ahyxccwylvytxcf; (2) zebrafish melanoma 10x Visium data (GEO repository: GSE159709); (3) human tonsil spatial CITE-seq data (https://www.10xgenomics.com/datasets/gene-protein-expression-library-of-human-tonsil-cytassist-ffpe-2-standard); (4) human HER2-positive breast tumor spatial transcriptomics data (https://github.com/almaan/her2st).

Details of the datasets analyzed in this paper were described in **Supplementary Table 1**.

## Software availability

An open-source implementation of the MorphLink algorithm can be downloaded from GitHub: https://github.com/jianhuupenn/MorphLink

## Life sciences reporting summary

Further information on experimental design is available in the Life Sciences Reporting Summary.

