## Supplementary Information for "MorphLink: Bridging Cell Morphological Behaviors and Molecular Dynamics in Multi-modal Spatial Omics"

**Supplementary Table 1. Image features extracted from each mask in MorphLink.**

| Feature name | Explanation |
|---|---|
| **Mask-level features** | |
| Area_of_1 | Area of white pixels in the mask |
| Area_of_1_ratio | Area of white pixels /total area of the mask |
| Dist_Trans_0_mean | Mean of the distance transformation matrix for black pixels |
| Dist_Trans_0_median | Median of the distance transformation matrix for black pixels |
| Dist_Trans_0_std | Standard deviation of the distance transformation matrix for black pixels |
| Dist_Trans_0_iqr | IQR deviation of the distance transformation matrix for black pixels |
| Dist_Trans_1_mean | Mean of the distance transformation matrix for white pixels |
| Dist_Trans_1_median | Median of the distance transformation matrix for white pixels |
| Dist_Trans_1_std | Standard deviation of the distance transformation matrix for white pixels |
| Dist_Trans_1_iqr | IQR deviation of the distance transformation matrix for white pixels |
| **Object-level features** | |
| area | Area of the object |
| bbox_area | Area of bounding box of the object |
| convex_area | Area of minimum convex polygon of the object |
| eccentricity | Eccentricity of the ellipse that has the same second-moments as the object |
| equivalent_diameter | The diameter of a circle with the same area as the object |
| extent | Ratio of pixels in the object to pixels in the total bounding box |
| filled_area | Area of the object with all the holes filled in |
| major_axis_length | The length of the major axis of the ellipse that has the same normalized second central moments as the object |
| minor_axis_length | The length of the minor axis of the ellipse that has the same normalized second central moments as the object |
| orientation | Counter-clockwise angle between the 0th axis (rows) and the major axis of the ellipse that has the same second moments as the object |
| perimeter | Perimeter of object which approximates the contour as a line through the centers of border pixels using a 4-connectivity |
| solidity | Ratio of pixels in the region to pixels of the convex hull image |
| hw_ratios | Ratio between major and minor axis |

Summary statistics for each object-level measurments, including mean, median, interquartile range (IQR), standard deviation, and quantiles (ranging from 0 to 1 in steps of 0.25), are compiled for each patch.

**Supplementary Table 2. Datasets analyzed in this paper.**

| Species | Tissue | Data source | Dataset dimensions | Protocol | Patch Size/pixels |
|---|---|---|---|---|---|
| Human | Bladder tumor | https://www.ncbi.nlm.nih.gov/geo/query/acc.cgi?acc=GSE246011 | 9,029 spots 18,085 genes | 10x Visium | 400 |
| Zebrafish | Embryo | Hunter *et al*.[1] (GSE159709) | 9,029 spots 18,085 genes | 10x Visium | 400 |
| Human | Tonsil | 10x Genomics (https://www.10xgenomics.com/datasets/gene-protein-expression-library-of-human-tonsil-cytassist-ffpe-2-standard) | 4,194 spots 18,060 genes 35 proteins | Spatial CITE-seq | 360 |
| Human | HER2-positive breast tumor | Andersson *et al*.[2] (https://github.com/almaan/her2st) | 295 spots 15,109 genes | Spatial Transcriptomics | 280 |

**Supplementary Table 3. Masks property summary for bladder cancer dataset.**

| Mask (Explanation) | % of patches contain this mask | Median % of area in patches contain this mask | Median RGB value | Mask examples |
|---|---|---|---|---|
| Mask1 (Nuclei) | 98.1% | 7.8% | 99, 33, 68 | 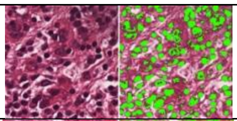 |
| Mask2 (CAF) | 99.0% | 21.6% | 148, 52, 89 | 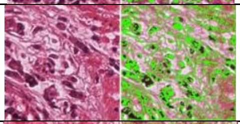 |
| Mask3 (Stroma1) | 99.0% | 14.5% | 188, 81, 122 | 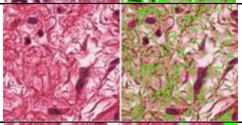 |
| Mask4 (Background1) | 95.7% | 22.3% | 241, 229, 232 | 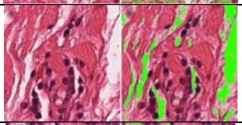 |
| Mask5 (Background2) | 97.6% | 11.3% | 240, 203, 217 | 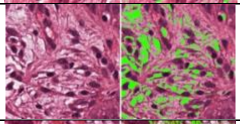 |
| Mask6 (Fiber1) | 98.9% | 19.5% | 207, 87, 124 | 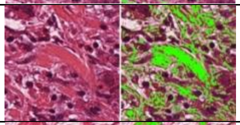 |
| Mask7 (Fiber2) | 98.8% | 23.5% | 177, 59, 92 | 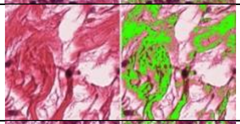 |
| Mask8 (Stroma2) | 98.4% | 10.2% | 230, 149, 179 | 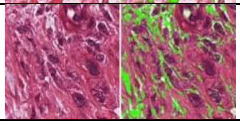 |

**Supplementary Table 4. Masks property summary for breast cancer dataset on sample H1.**

| Mask (Explanation) | % of patches contain this mask | Median % of area in patches contain this mask | Median RGB value | Mask examples |
|---|---|---|---|---|
| Mask1 (Background) | 99.8% | 32.5% | 227, 205, 226 | 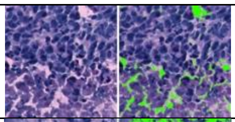 |
| Mask2 (Stroma1) | 100.0% | 31.6% | 174, 148, 190 | 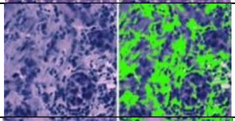 |
| Mask3 (Stroma2) | 100.0% | 14.9% | 126, 114, 168 | 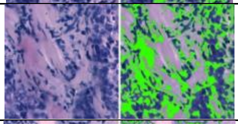 |
| Mask4 (Nuclei) | 99.5% | 13.9% | 50, 59, 120 | 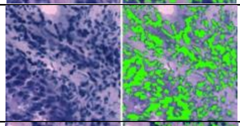 |
| Mask5 (Nuclei boundary) | 100.0% | 8.3% | 90, 89, 148 | 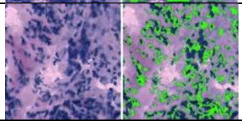 |

**Supplementary Table 5. Masks property summary for zebrafish melanoma dataset.**

| Mask (Explanation) | % of patches contain this mask | Median % of area in patches contain this mask | Median RGB value | Mask examples |
|---|---|---|---|---|
| Mask1 (Melanoma 1) | 94.7% | 27.8% | 124, 80, 153 | 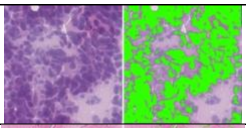 |
| Mask2 (Nuclei boundary) | 92.7% | 6.0% | 181, 123, 176 | 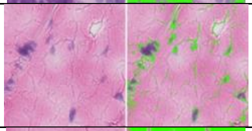 |
| Mask3 (Fiber bundles 1) | 77.3% | 47.6% | 219, 144, 189 | 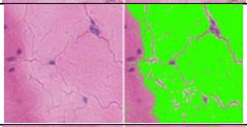 |
| Mask4 (Fiber bundles at cell boundary) | 88.7% | 9.2% | 194, 107, 162 | 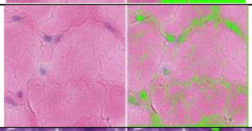 |
| Mask5 (Melanoma 2) | 19.2% | 4.2% | 79, 34, 68103 | 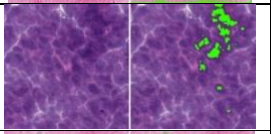 |
| Mask6 (Inter-cellular space) | 45.9% | 9.9% | 223, 216, 222 | 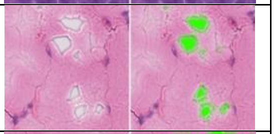 |
| Mask7 (Fiber bundles 2) | 77.4% | 8.3% | 224, 171, 205 | 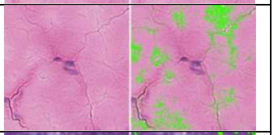 |
| Mask8 (Melanoma 3) | 53.7% | 12.8% | 104, 56, 133 | 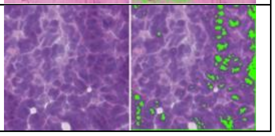 |

**Supplementary Table 5. Masks property summary for human tonsil dataset.**

| Mask (Explanation) | % of patches contain this mask | Median % of area in patches contain this mask | Median RGB value | Mask examples |
|---|---|---|---|---|
| Mask1 (Background1) | 91.2% | 11.4% | 232, 217, 235 | 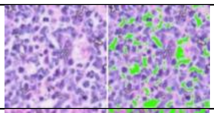 |
| Mask2 (Background2) | 99.6% | 20.4% | 222, 191, 233 | 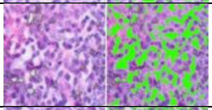 |
| Mask3 (Stroma1) | 98.1% | 21.9% | 193, 157, 223 | 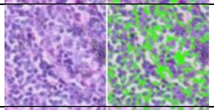 |
| Mask4 (Nuclei light1) | 97.8% | 20.1% | 164, 129, 206 | 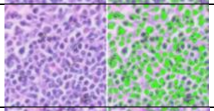 |
| Mask5 (Nuclei light2) | 97.7% | 13.2% | 132, 98, 183 | 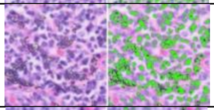 |
| Mask6 (Nuclei dark) | 95.4% | 5.3% | 115, 82, 166 | 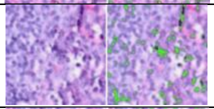 |
| Mask7 (Stroma1) | 76.1% | 9.9% | 209, 152, 221 | 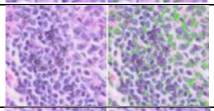 |
| Mask8 (Stroma2) | 43.5% | 7.9% | 202, 126, 206 | 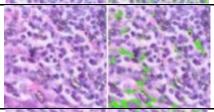 |
| Mask9 (Fiber) | 42.3% | 9.9% | 41, 109, 168 | 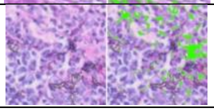 |

**Supplementary Fig. 1.** SVGs associated with antigen-presenting and tumor proliferation are enriched in Tumor Subregion 1 in the bladder cancer data.

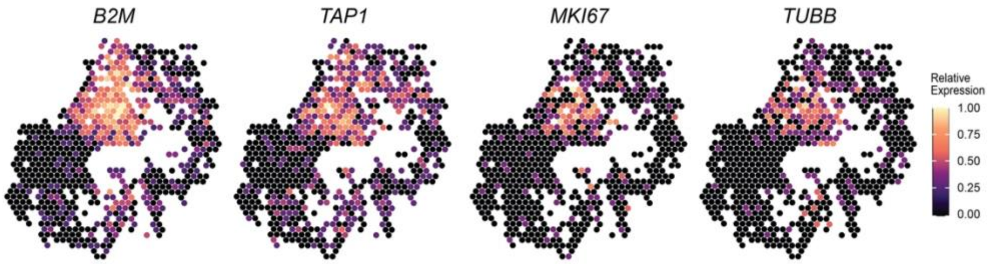

**Supplementary Fig. 2.** All eight masks and detected objects for spots 1 and 2 in the bladder cancer data.

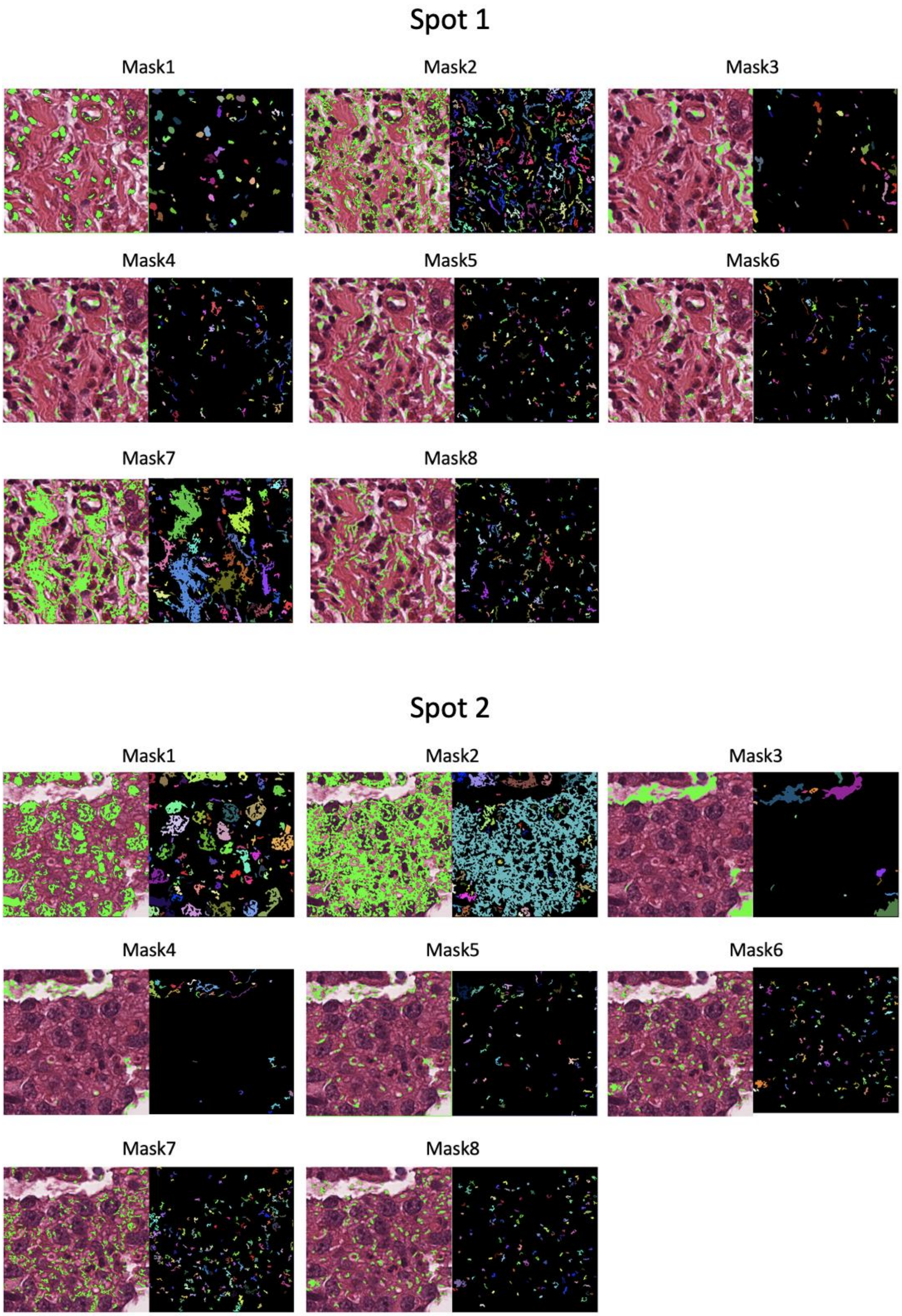

**Supplementary Fig. 3.** Expression of genes that involved in regulating transcriptional activity and chromatin remodeling in the bladder cancer data.

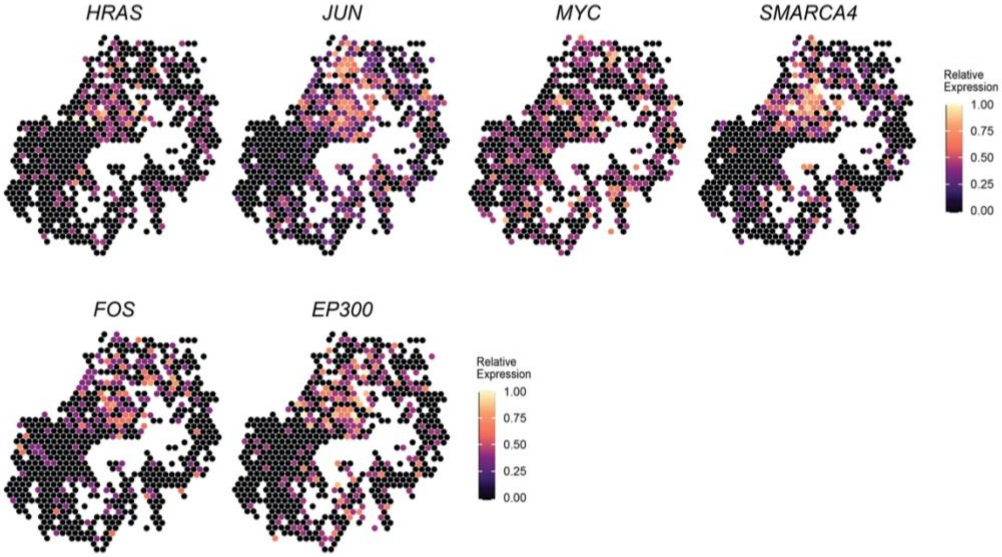

**Supplementary Fig. 4**. The distribution of CPSIs between *IGHM*, *MS4A1* and all lymphocyte nuclei features in the bladder cancer data.

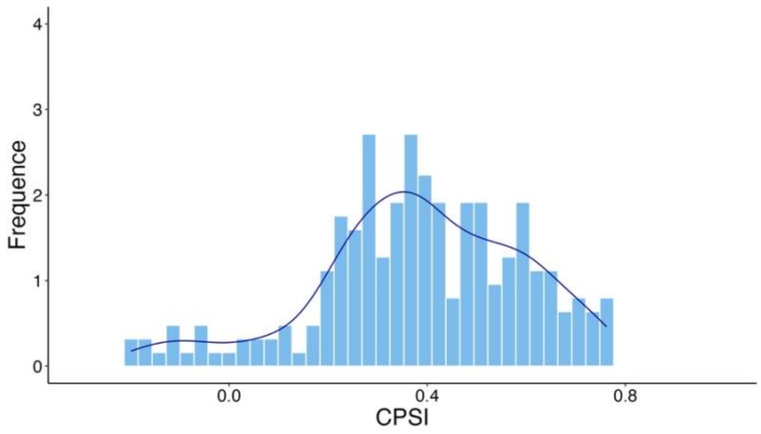

**Supplementary Fig. 5**. Expression of collagen genes pattern in tumor region in A1, G2 and H1 samples.

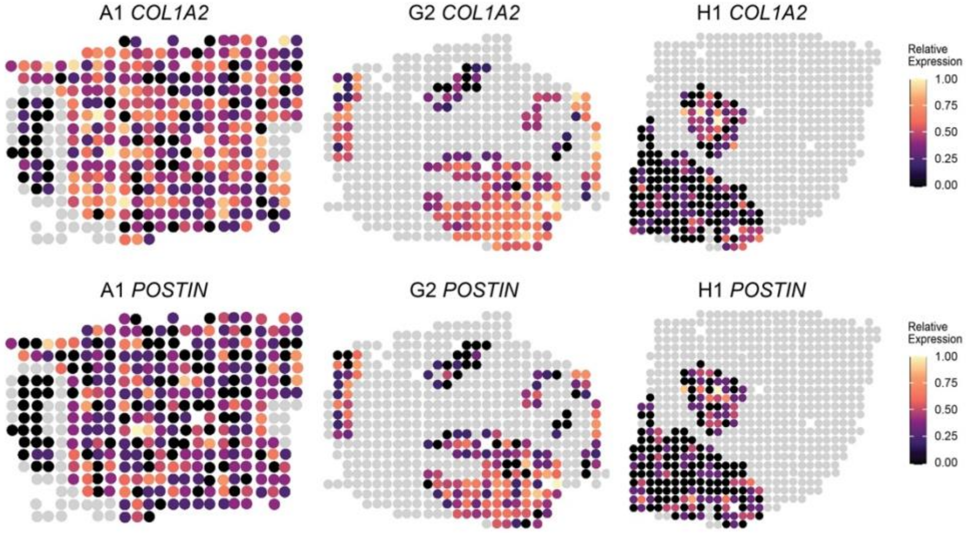

**Supplementary Fig. 6**. Distibution of CPSIs between *COL1A1* and stroma morphology features in each sample, CPSI between *COL1A1* and selected feature equals 0.454, 0.379, 0.165 in A1, G2, H1 sample.

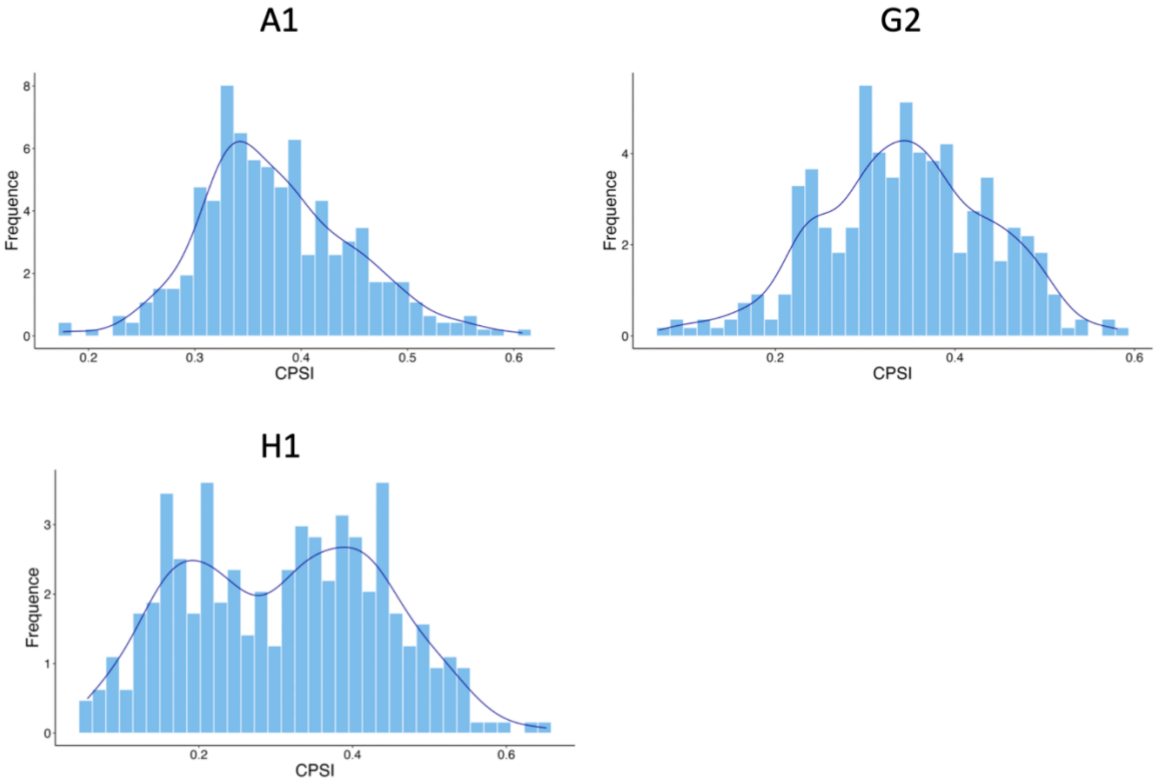

**Supplementary Fig. 7** A visual illustration depicting of image feaure 1 in zebrafish data. The values are median image feature values for spot stratified by *RPL15*'s expression level, grouped into quantiles from 0 to 1 with step of 0.25.

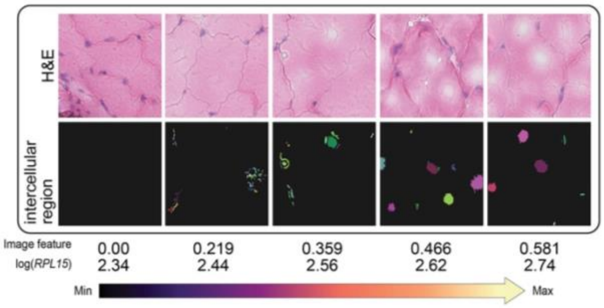

**Supplementary Fig. 8**. Tumor/muscle enriched genes in the zebrafish dataset.

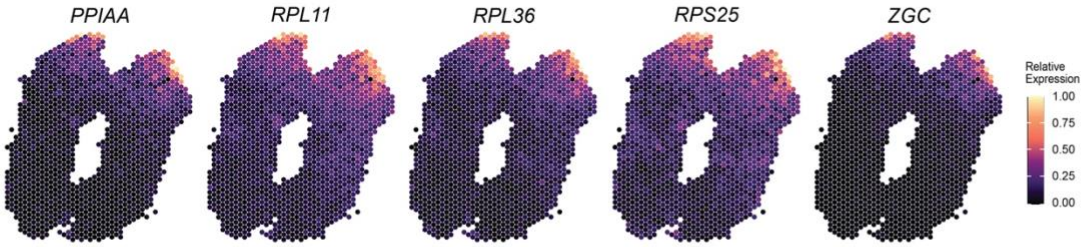

**Supplementary Fig. 9**. Distibution of CPSIs between muscle/tumor interface enriched genes and fiber morphology features in zebrafish data. CPSI between the selected image feature(area of fiber bundles, quantile 0.75) with each genes are *RPL5B* (0.757), *TUBA8L* (0.743), *PPIAA* (0.747), *ZGC* (0.724), *RPL11* (0.718), *RPL41* (0.689), *RPS25* (0.701), *RPL36* (0.726).

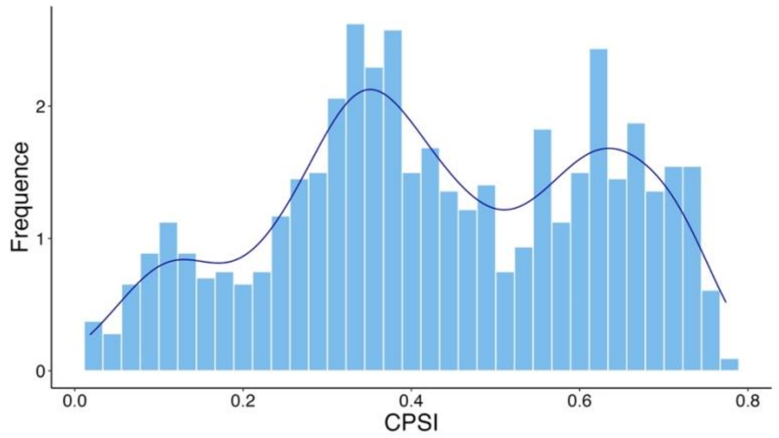

**Supplementary Note 1: The features from MorphLink exhibit greater robustness to batch effects compared to HIPT features.**

To evaluate the robustness of extracted image features to batch effects, we employed MorphLink and HIPT[3] on the HER2+ breast cancer dataset, consisting of eight annotated sections from different samples. Based on extracted image features, we first performed PCA to reduce the dimension and retained the top 50 PCs for UMAP visualization. From the UMAP plot, each dot represents an image feature extracted from a single section, with its color indicating the sample source (shown in **Supplementary Fig. 10**). The image features from MorphLink are more mixed across samples, while those from HIPT are aggregated into clusters by their sample sources. The distinction in patterns arises from the interpretability and transparency nature of the extracted features from MorphLink. Their context-based meanings enable them to be more comparable across different sections and samples, in contrast to those with unclear implications from deep neural networks. Therefore, we conclude that MorphLink features are scalable to multi-sample data and show great robustness to batch effects.

**Supplementary Fig. 10.** UMAP of spots from eight samples using features from MorphLink and HIPT.

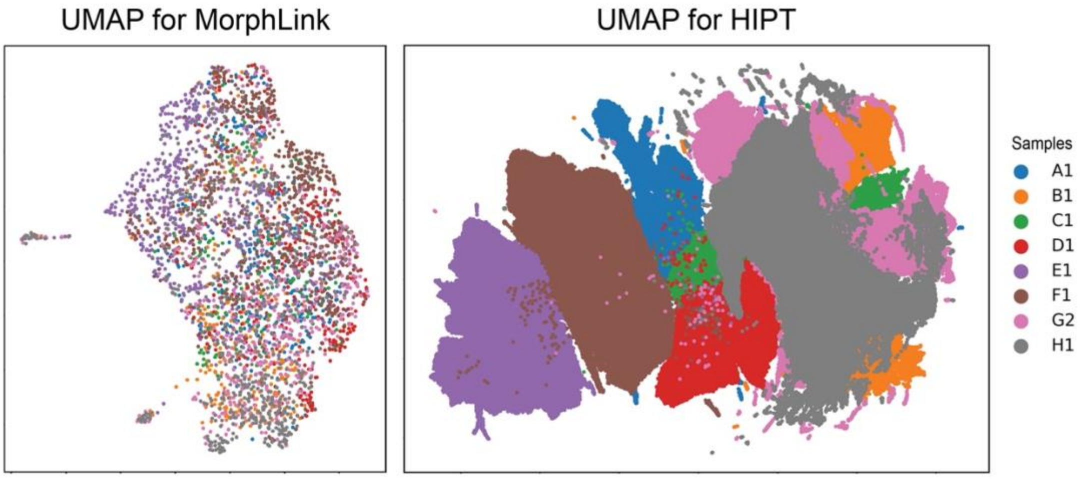

**Supplementary Note 2: CPSI demonstrates superior performance in quantifying spatial pattern similarity across different simulated scenarios.** To compare the capability of CPSI with other metrics in identifying coherent spatial patterns, we simulated five features with distinct spatial expression patterns as the evaluation dataset. In this evaluation, we selected feature 0 as the target feature and used various metrics to quantify the spatial pattern similarity between feature 1 and features 2 to 5. As shown in **Supplementary Fig. 11a**, feature 1 exhibits identical values in the upper-left triangle and a gradient of decreasing values along the x and y axes in the lower-right triangle area. Features 2 and 3 are considered positive samples due to their similar patterns to feature 1, with identical values in the upper-left triangle and decreasing values along the x and y axes at different rates in the lower-right triangle area. In contrast, features 4 and 5 are considered negative samples, as feature 4 shows a random pattern and feature 5 exhibits an opposite increasing gradient along the x and y axes in the lower-right triangle area. The bar plot in **Supplementary Fig. 11b** displays the spatial similarity quantification results from different metrics. Both positive samples, feature 2 and 3, have the highest positive CPSI values, close to 1, indicating perfect pattern similarity. Feature 4, with random patterns, shows a CPSI close to 0, implying no pattern association with feature 1. Feature 5 has a negative CPSI of -0.25, suggesting opposite spatial patterns. Correlation and SSIM show similar quantification results. They successfully identify feature 2 as having a similar pattern with high values and feature 4 as having an irrelevant pattern, indicated by values close to 0. However, both correlation and SSIM mistakenly identify feature 3 as having a dissimilar pattern with negative values and feature 5 as having a relatively similar pattern with positive values. These mistakes suggest that these two metrics tend to average global spatial patterns but overlook local distinct gradients. RMSE identifies the similar pattern of feature 3 with a value of 0.5 but generally fails the quantification task for other features.

**Supplementary Fig. 11. Benchmark CPSI with other metrics using simulated features. a.** Simulated expression patterns of 5 features. **b.** Barplot for CPSI, correlation, SSIM, and RMSE calculated between Feature 1 and Feature 2 – 5.

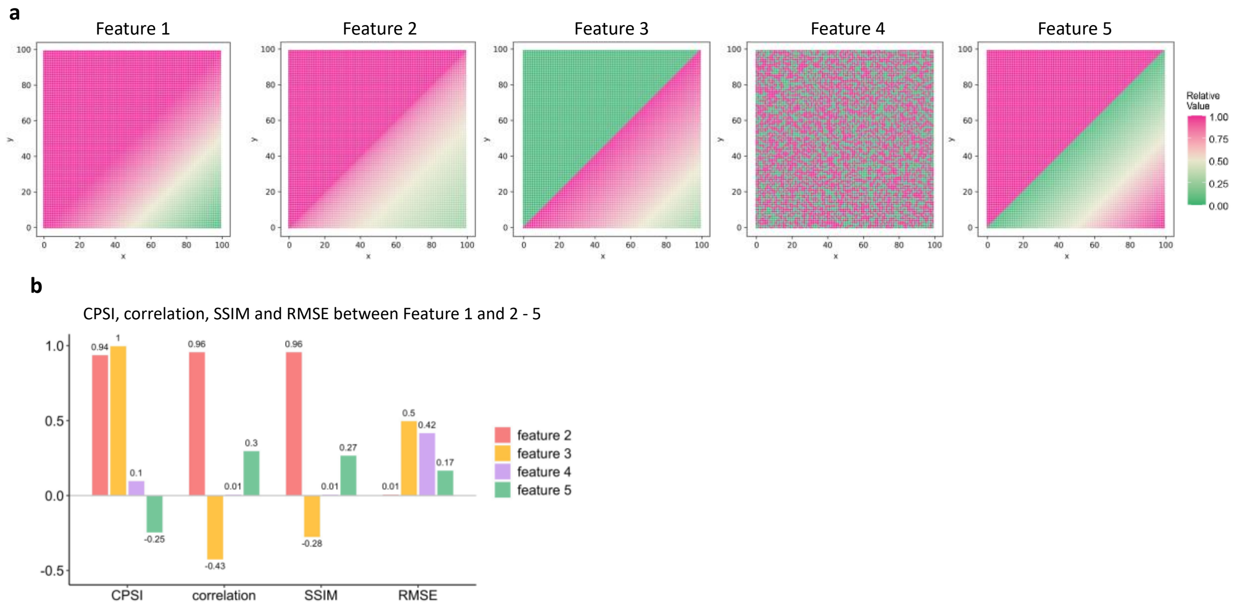

**Supplementary Note 3: Morphology-molecular linkage detection in tri-modality data**
To showcase MorphLink's capability with multi-modal spatial omics data, we applied it to a tri-modality CITE-seq dataset from human tonsil tissue, which included H&E imaging, spatial transcriptomics, and proteomics. As highlighted in **Supplementary Fig. 12a**, the tonsil is characterized by its secondary lymphoid follicle structure, which is crucial for initiating adaptive immunity[4]. These follicles display significant immune environment, primarily featuring a germinal center surrounded by a mantle zone. This structure can be distinguished from other forms through clustering analysis using proteomics data, identified as clusters 3 and 7 in **Supplementary Fig. 12b**. PD-1 is an inhibitory immune checkpoint receptor, known for its expression on T cells. It functions as an inhibitory receptor to modulate immune responses and maintain self-tolerance by limiting autoimmune reactions. Its high abundance in germinal centers (**Supplementary Fig. 12c**), is important in regulating follicular helper T cells (Tfh), which play a critical role in B cell affinity maturation and selection. Beginning with this protein, we showcase that MorphLink can identify a tri-modality linkage of protein-gene-morphology that elucidates the unique immune processes in the germinal center and mantle zone. The gene *PDCD1* exhibited the 2$^{nd}$ highest CPSI score of 0.652, as detailed in **Supplementary Fig. 12d**. Moreover, *TIGIT*, as illustrated in **Supplementary Fig. 12e**, was identified as the top gene with a pattern similar to PD-1, achieving the highest CPSI score of 0.663. Prior research[5, 6] has shown that *TIGIT* and *PDCD1* are often co-expressed in Tfh and NK cells in various tumor types. Additional studies have demonstrated the potential benefits of co-inhibition of TIGIT and PD-1/PD-L1 in enhancing anti-tumor immunity and improving treatment outcomes in several cancer types[5]. Although *TIGIT*'s protein abundance is not directly measured, this protein-gene linkage suggests a synergistic function between *PDCD1* and *TIGIT* in regulating Tfh at the germinal center. This regulation influences the process of B cell affinity maturation and selection, driving various immune functions in the germinal center and mantle zone.

Besides gene expression, MorphLink identified an image feature (**Supplementary Fig. 12f**) spatially aligned with PD-1 (protein) (CPSI = 0.545). From **Supplementary Table 5**, we can find that this feature from mask 4 quantifies the distribution of lymphocytes with dark, dense chromatin. As illustrated in **Supplementary Fig. 12g**, low values in this feature indicates a high concentration of small, resting B cells with dense chromatin in the mantle zone. In contrast, a high value of this feature identified less densely stained lymphocytes within the germinal center. These lymphocytes are larger in size and exhibit more open chromatin, reflecting their more active roles in responding to antigens inside the germinal center. By utilizing tri-modality data, MorphLink identifies co-expression patterns between the PD-1 protein and the genes *PDCD1* and *TIGIT*, and further correlates these molecular signatures with distinctive B cell morphologies that reflect their functional states. This holistic approach enables a deeper understanding of unique biological processes and the dynamic interplay between cellular morphology and molecular activity in immune regulation.

**Supplementary Fig. 12. MorphLink identifies associations among morphology, gene expression, and protein abundance in a human tonsil spatial CITE-seq dataset. a.** The H&E image of human tonsil tissue, and a zoomed-in view of the secondary lymphoid follicle structure. **b.** Spatial domains detected using protein abundance. **c.** Abundance of PD-1. **d.** Expression pattern of *PDCD1*. **e.** Expression pattern of *TIGIT*. **f.** A morphological feature that quantifies the IQR of distances between lymphocyte pixels has the highest CPSI with PDCD1 (protein). **g.** A visual illustration depicting changes in the concentration of lymphocytic nuclei associated with PDCD1 abundance in both the H&E image and the segmented mask.

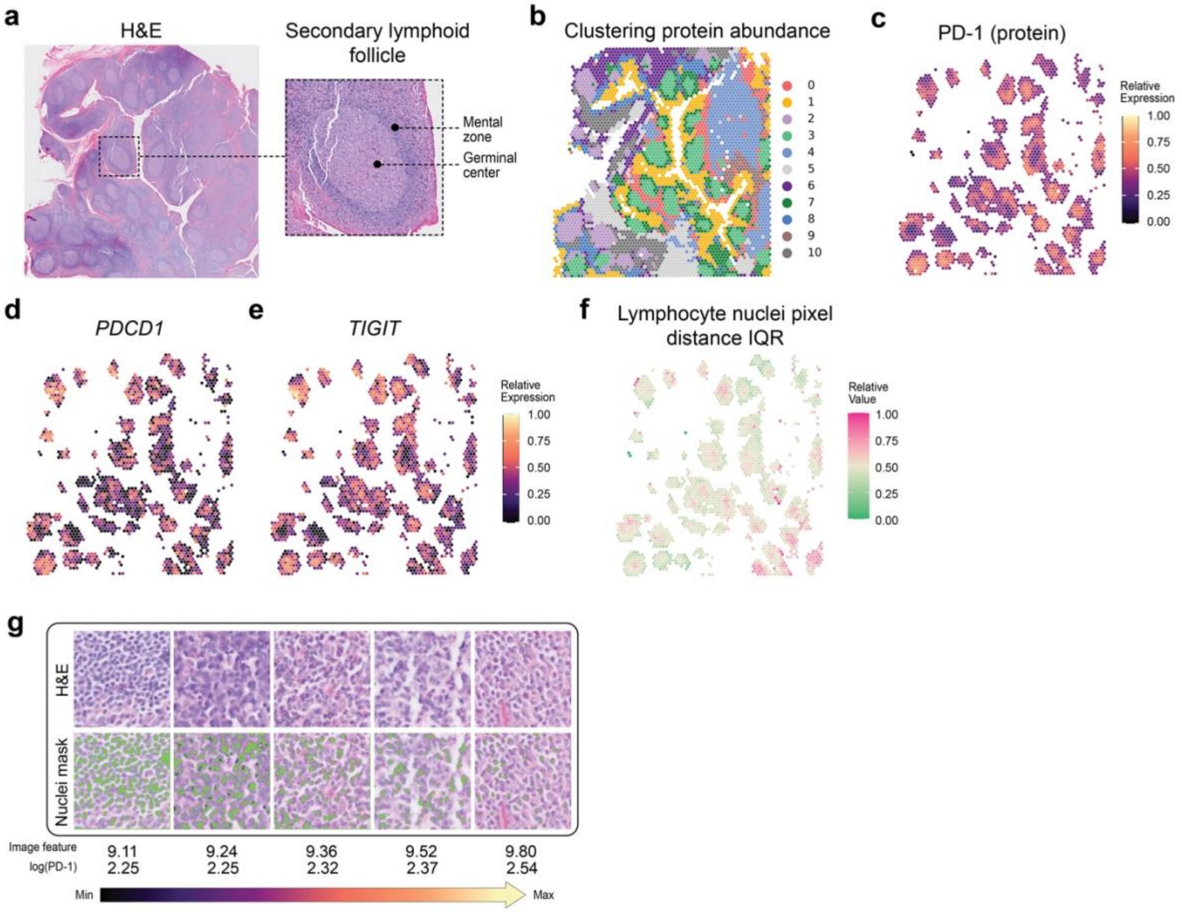
